# Mitochondrial protein import couples proteostasis failure to mitochondrial permeabilization

**DOI:** 10.64898/2026.09.03.749044

**Authors:** Zhiqi Sun, Hauke Holthusen, Sara Berndl, Antonia Behnsen, Sophia J. Schwojer, Giulia Gobbato, Grace Sorensen, Bettina Warscheid, Stephan A. Sieber, F. Ulrich Hartl, Veit Hornung

## Abstract

Proteostasis failure is a hallmark of stress and disease, yet how it compromises mitochondrial integrity remains unclear. Here, we identify mitochondrial protein import as a critical pathway linking proteostasis failure to mitochondrial injury. We show that Raptinal, previously characterized as a rapid inducer of apoptosis, impairs the folding of newly synthesized proteins rather than directly disrupting mitochondrial membranes. The resulting proteotoxic stress drives mitochondrial outer membrane permeabilization and intrinsic apoptosis independently of BCL-2 family pore-forming proteins. VBIT4, a compound commonly used to maintain mitochondrial integrity, inhibited this pathway, and chemical proteomics with a photoaffinity analogue implicated the TIM23 import machinery. Genetic or pharmacological inhibition of the TIM23–PAM axis suppressed mitochondrial permeabilization without affecting canonical BAX–BAK-dependent apoptosis. These findings establish that mitochondrial protein import couples translation-associated proteotoxic stress to mitochondrial injury and identify regulation of import flux as a determinant of mitochondrial integrity during proteostasis failure.

## Introduction

Mitochondria support cellular bioenergetics and function as central signaling hubs that regulate cell fate. In intrinsic apoptosis, mitochondrial outer membrane permeabilization (MOMP) allows intermembrane-space proteins, including cytochrome c and SMAC, to enter the cytosol, where cytochrome c promotes APAF1 apoptosome assembly and caspase-9 activation^1, 2^. This canonical form of MOMP is executed by the BCL-2 family effectors BAX and BAK and is controlled by the balance between pro- and anti-apoptotic BCL-2 family proteins^3, 4^. However, mitochondria can also undergo limited or non-canonical permeabilization that releases mitochondrial damage-associated molecular patterns (DAMPs), including mitochondrial DNA (mtDNA), without immediate apoptotic cell death^5,6^. In addition to sublethal BAX/BAK-dependent permeabilization^6, 7^, oligomerization of voltage-dependent anion channels (VDACs) in the outer mitochondrial membrane has been proposed as a mechanism that enables mtDNA release into the cytosol^8^. Mitochondria-derived DAMPs contribute to inflammatory signaling in senescence^6, 9^, neurodegeneration^10, 11^, and aging^12^, yet the cellular stresses that initiate non-canonical mitochondrial permeabilization remain poorly understood.

Proteostasis failure frequently accompanies mitochondrial dysfunction in neurodegeneration and aging^13–15^. One vulnerable point in the proteostasis network is the folding of newly synthesized proteins, which occurs as proteins are sorted to their final cellular destinations^16^. This vulnerability may be especially pronounced for mitochondria, because most mitochondrial proteins are synthesized in the cytosol and imported across mitochondrial membranes^17^. Substrates of the TOM–TIM23 import pathway, including most matrix and many presequence-containing inner-membrane proteins, must remain at least partially unfolded during translocation and complete folding only after import, creating a constant burden on mitochondrial proteostasis. Together, these features raise the possibility that mitochondrial protein import provides a route through which proteostasis failure can compromise mitochondrial integrity.

Raptinal is a small molecule widely used to induce rapid apoptosis^18^ and previously shown to trigger mitochondrial permeabilization independently of BCL-2 family pore-forming proteins^19^. We unexpectedly found that Raptinal acutely impairs productive folding of newly synthesized proteins, thereby inducing translation-associated proteotoxic stress upstream of mitochondrial permeabilization. We further established mitochondrial protein import as a critical vulnerability through which proteostasis failure compromises mitochondrial integrity.

## Results

### Raptinal induces translation-associated proteostasis stress upstream of mitochondrial damage

To access non-canonical mitochondrial permeabilization independently of canonical apoptosis, we first confirmed that Raptinal-induced death proceeds without BAX and BAK^19^. In pooled HEK293 knockouts, Raptinal-induced apoptosis was independent of caspase-8 and BAX/BAK but dependent on caspase-9 (Supplementary Fig. 1a,b), placing mitochondrial damage upstream of caspase activation and distinguishing this pathway from classical BAX/BAK-dependent apoptosis. This non-canonical route is not restricted to a single model, as Raptinal induced rapid, dose-dependent caspase-3/7 activation at concentrations above 2.5 µM within 2 hours in HEK293T, HEK293, and HeLa cells and across multiple other cell lines^18^ (Supplementary Fig. 1c). In contrast to HeLa cells, HEK293T and HEK293 cells were resistant to BH3-mimetic-induced apoptosis^20, 21^ (Supplementary Fig. 1d). This differential sensitivity provided an experimental system to examine Raptinal-induced mitochondrial permeabilization under conditions in which canonical BAX/BAK-dependent MOMP is inefficient. We therefore used HEK293 cells for mechanistic analyses and HeLa cells as a comparator for canonical BH3-mimetic-induced apoptosis.

To directly assess MOMP independently of downstream caspase activity, we titrated Raptinal in CASP9^KO^ HEK293 cells and isolated cytosolic fractions by mild digitonin permeabilization. Under basal conditions, neither cytochrome c nor SMAC was detected in the cytosol, indicating preservation of mitochondrial integrity during extraction. Upon Raptinal treatment, cytochrome c release was observed at concentrations above 2.5 µM, whereas SMAC release was detectable at lower concentrations (Supplementary Fig. 1e), suggesting differential sensitivity of intermembrane space proteins to Raptinal-induced permeabilization. Notably, Raptinal failed to permeabilize isolated mitochondria even at 50 µM, whereas digitonin efficiently disrupted the outer membrane at concentrations above 0.025% (w/v) (Supplementary Fig. 1f). Thus, Raptinal does not directly compromise mitochondrial membranes but instead requires cellular context, implying the involvement of upstream cellular processes.

To identify upstream pathways while minimizing confounding effects of caspase activation, we performed RNA-seq in CASP9^KO^ HEK293 cells (Supplementary Table 1). Pathway analysis revealed coordinated activation of stress-response programs by Raptinal, prominently including an ATF4-associated integrated stress response (ISR)^22^ and an HSF1-driven heat-shock response^23^ (Fig. 1a). Among the most strongly induced transcripts were canonical ATF4 targets (DDIT3, SLC7A11, TRIB3) and heat-shock genes (DNAJB1, BAG3, HSPA6) (Fig. 1b,c). qPCR analysis confirmed robust induction of these targets (Fig. 1d). Inhibition of protein translation with cycloheximide (CHX) suppressed induction of DNAJB1, BAG3, and HSPA6 and markedly reduced DDIT3 expression, indicating that activation of the heat-shock response, and to a lesser extent the ISR, depends on ongoing translation. By contrast, treatment with ISRIB^24^, a small-molecule inhibitor of the ISR, attenuated DDIT3 induction while further enhancing heat-shock gene expression. These data indicate that Raptinal induces a translation-associated proteostasis stress response characterized by concurrent activation of HSF1- and ATF4-driven pathways.

**Figure 1.**
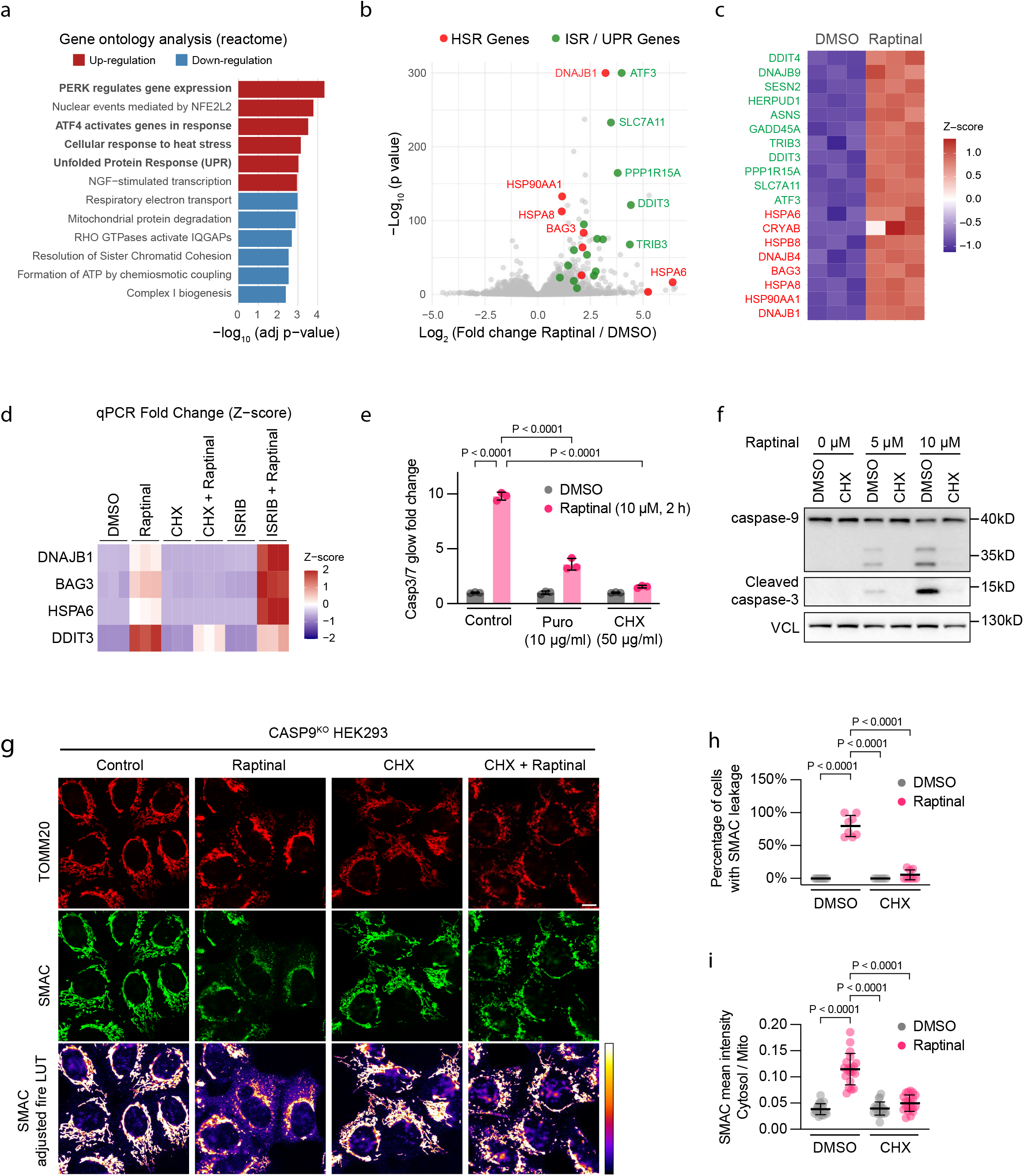
Raptinal induces translation-associated proteostasis stress upstream of mitochondrial damage. **(a)** CASP9^KO^ HEK293 cells were treated with DMSO or Raptinal (10 μM) for 6 h, followed by RNA- seq and differential expression analysis using DESeq2. Reactome pathway enrichment was performed on 113 upregulated genes (Wald statistic > 10) and 135 downregulated genes (Wald statistic < −6). Pathways are ranked by adjusted p values. **(b)** Volcano plot of Raptinal-induced transcriptional changes, highlighting strong induction of ISR/UPR- and HSR-associated transcripts (n = 3). **(c)** Heatmap showing z-scored variance-stabilizing transformation (VST)-normalized expression of ISR/UPR- and HSR-associated transcripts highlighted in **(b)**. **(d)** CASP9^KO^ HEK293 cells were treated with DMSO or 10 μM Raptinal, in the presence or absence of CHX (50 µg/mL) or ISRIB (200 nM) for 6 h. Expression of selected ATF4- and HSF1-associated genes was analyzed by RT-qPCR, normalized to GAPDH. Fold changes were standardized within each gene and displayed as z-scores. n = 3 independent experiments. **(e)** Wild-type HEK293 cells were pretreated with vehicle, puromycin (10 μg/mL), or CHX (50 μg/mL) for 1 h and then treated with 10 μM Raptinal for 2 h. Caspase-3/7 activity was measured by Caspase- Glo 3/7, normalized to DMSO-treated control and shown as mean ± SD; n = 3 independently treated wells, representative of 3 independent experiments; two-way ANOVA with Tukey’s correction. **(f)** Wild-type HEK293 cells were pretreated with vehicle or CHX (50 μg/mL) for 1 h followed by 5 or 10 μM Raptinal for 2 h. Caspase-9 processing and caspase-3 cleavage were analyzed by western blotting. Vinculin (VCL), loading control. Representative of 2 independent experiments. **(g)** CASP9^KO^ HEK293 cells were pretreated with vehicle or CHX (50 μg/mL) for 1 h followed by DMSO or Raptinal (10 μM) for 2 h. Cells were fixed and immunostained for TOMM20 and SMAC. Representative confocal images are shown with TOMM20 in red and SMAC in green or contrast- enhanced Fire LUT. Scale bar, 10 μm. **(h)** Percentage of cells with increased cytosolic SMAC signal after treatment as in **(g)**. Each point represents one field; bars indicate mean ± SD; n = 6-8 fields per condition pooled from 3 independent experiments; one-way ANOVA with Tukey’s correction. **(i)** Cytosolic-to-mitochondrial SMAC intensity ratio in cells treated as in **(g)**. Data are mean ± SD; n = 20 cells per condition pooled from 3 independent experiments; one-way ANOVA with Tukey’s correction.

We next asked whether this proteostasis stress is functionally linked to mitochondrial damage. Inhibition of the ISR with ISRIB had only a minor effect on Raptinal-induced apoptosis (Supplementary Fig. 2a). In contrast, inhibition of cytosolic translation with CHX markedly suppressed apoptosis in HEK293 cells (Fig. 1e), an effect partially recapitulated by puromycin (Fig. 1e). Consistently, CHX blocked Raptinal-induced cleavage of caspase-9 and caspase-3 (Fig. 1f), indicating that translation is required upstream of caspase activation. This requirement was not cell-type specific, as CHX similarly inhibited Raptinal-induced apoptosis in HeLa cells (Supplementary Fig. 2b). Notably, under the same conditions, CHX potentiated BH3-mimetic-induced apoptosis (Supplementary Fig. 2c), likely due to rapid turnover of the anti-apoptotic protein MCL1^25^, underscoring the distinct mechanistic basis of Raptinal-induced cell death.

To directly examine whether translation is required for MOMP, we monitored SMAC localization by immunofluorescence in CASP9^KO^ HEK293 cells. Under basal conditions, SMAC colocalized with TOMM20 within an interconnected tubular mitochondrial network. Raptinal treatment induced pronounced mitochondrial fragmentation and perinuclear clustering, accompanied by increased cytosolic SMAC signal (Fig. 1g). Quantitative analysis revealed that approximately 80% of cells exhibited a ∼3-fold increase in cytosolic SMAC intensity (Fig. 1h,i). However, a substantial fraction of SMAC remained mitochondrial, indicating incomplete permeabilization. This contrasts with the rapid and near-complete SMAC release observed during canonical BAX/BAK-mediated apoptosis^26^, suggesting that Raptinal induces a partial or non-canonical form of MOMP. Importantly, inhibition of translation with CHX abolished SMAC release (Fig. 1g-i) and partially restored mitochondrial network integrity (Supplementary Fig. 2d).

Collectively, these findings establish that Raptinal induces a translation-associated proteostasis stress that is required for mitochondrial outer membrane permeabilization.

### Direct impairment of productive protein folding underlies Raptinal-induced proteotoxic stress

To directly assess the effect of Raptinal on cellular proteostasis, we inducibly expressed HA-tagged wild-type firefly luciferase (Fluc-WT) in CASP9^KO^ HEK293 cells (Fig. 2a). Firefly luciferase folding, which can be quantified by enzymatic activity, is highly sensitive to cellular proteostasis^27^. We normalized luciferase activity to reporter protein abundance to derive specific activity as a proxy for folding efficiency. To minimize secondary effects from mitochondrial permeabilization, we treated cells with sublethal Raptinal concentrations below 1.25 µM during reporter induction. Raptinal reduced the specific activities of Fluc-WT in a dose-dependent manner, indicating impaired folding of newly synthesized Fluc reporters (Fig. 2b,c). By contrast, a 10-minute Raptinal pulse after luciferase induction did not reduce activity, arguing against assay interference or direct inhibition of the mature enzyme (Supplementary Fig. 3a,b).

**Figure 2.**
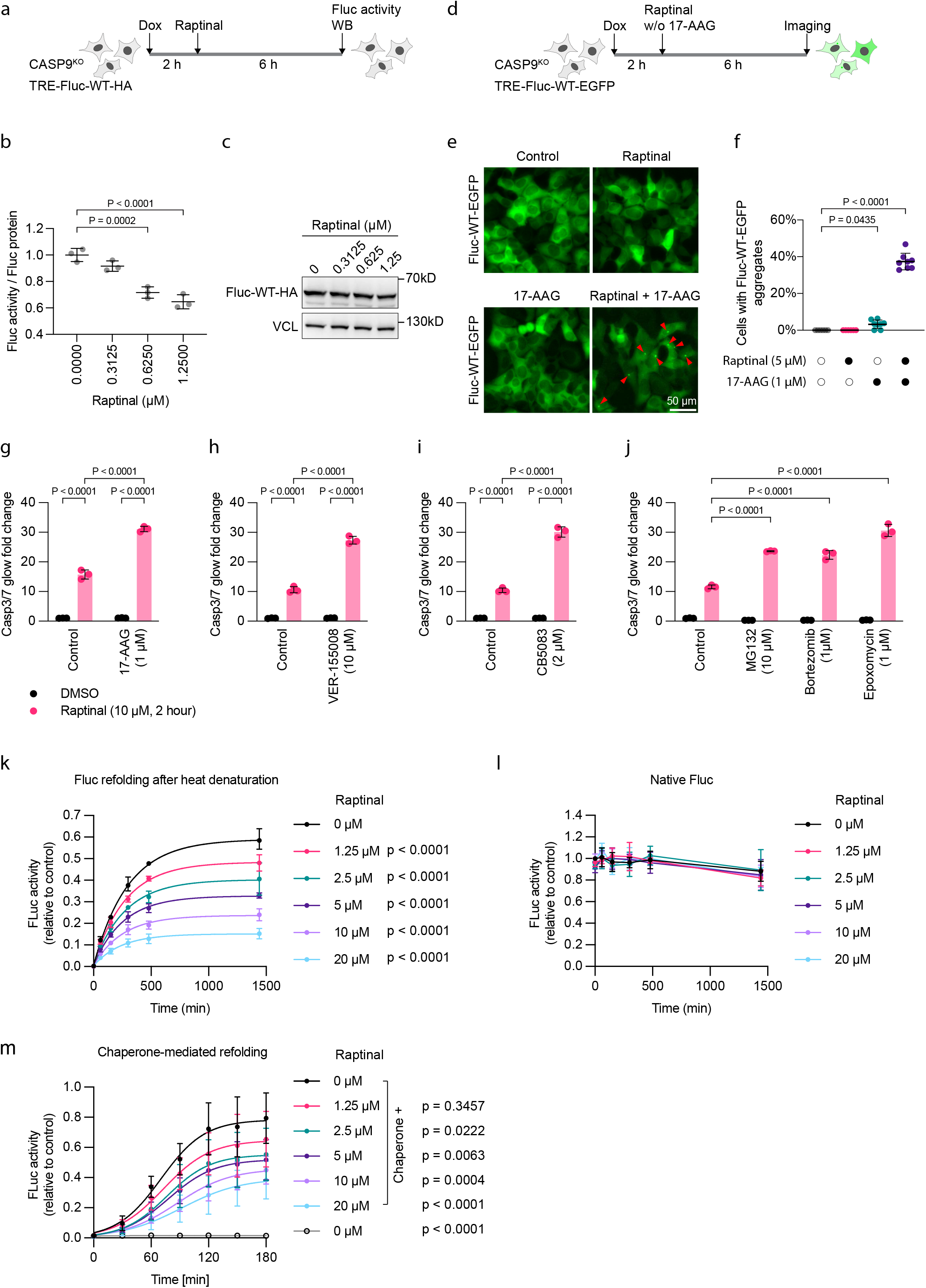
Direct impairment of productive protein folding underlies Raptinal-induced proteotoxic stress. **(a)** Experimental design for **(b)** and **(c)**. CASP9KO HEK293 cells inducibly expressing HA-tagged Fluc-WT were induced with doxycycline (Dox; 200 ng/mL, 2 h) and treated with Raptinal for 6 h. Luciferase activity and reporter abundance were measured. **(b)** Specific activity of firefly luciferase reporter from the experiment described in **(a)**, calculated by normalizing luciferase activity to reporter abundance. Data are mean ± SD; n = 3 independently treated wells, representative of 2 independent experiments. Statistical significance relative to Raptinal-free condition was calculated by one-way ANOVA with Dunnett’s correction. **(c)** HA-tagged firefly luciferase reporter abundance in cells treated as in **(a)**, analyzed by immunoblotting, representative of 2 independent experiments. VCL, loading control. **(d)** Experimental design for **(e)** and **(f)**. CASP9^KO^ HEK293 cells inducibly expressing EGFP-tagged wild-type firefly luciferase (TRE-Fluc-WT-EGFP) were treated with doxycycline (Dox; 200 ng/mL) for 2 h, followed by Raptinal (5 μM), 17-AAG (1 μM), or both for 6 h. Fluc-WT-EGFP was imaged by epifluorescence microscopy. **(e)** Representative epifluorescence images of Fluc-WT-EGFP in cells treated as in **(d)**. Fluc-WT-EGFP aggregates appear as bright intracellular foci and are indicated by red arrowheads. Scale bar, 50 μm. **(f)** Quantification of the percentage of cells with Fluc-WT-EGFP aggregates from images shown in **(e)**. Data are mean ± SD. n = 6-8 fields per condition pooled from 3 independent experiments. Statistical significance relative to control condition was calculated by one-way ANOVA with Dunnett’s correction. **(g–j)** Wild-type HEK293 cells were pretreated for 1 h with vehicle or the indicated proteostasis inhibitors: **(g)** HSP90 inhibitor 17-AAG (1 μM), **(h)** HSP70 inhibitor VER-155008 (10 μM), **(i)** p97/VCP inhibitor CB5083 (2 μM), or **(j)** proteasome inhibitors MG132 (10 μM), bortezomib (1 μM), or epoxomicin (1 μM), followed by treatment with Raptinal (10 μM) for 2 h. Caspase-3/7 activity was normalized to DMSO-treated control. Data are mean ± SD; n = 3 independently treated wells, representative of 3 independent experiments; two-way ANOVA with Tukey’s correction. **(k)** Recombinant Fluc was heat-denatured at 42 °C for 10 min with the indicated Raptinal concentrations and shifted to 25 °C for spontaneous refolding. Fluc activity was measured over time and normalized to native Fluc activity at each time point. Data are mean ± SD; n = 3 independent replicates. Data were fitted with one-phase association model. Statistical significance relative to the Raptinal-free condition at the end point was calculated by two-way ANOVA with Dunnett’s correction. **(l)** Native recombinant Fluc was incubated with the indicated Raptinal concentrations at 25°C for 24 h. Fluc activity was normalized to untreated native Fluc at 0 h. Data are mean ± SD; n = 3 independent replicates. **(m)** Recombinant Fluc was heat-denatured with the HSP70 chaperone system and the indicated Raptinal concentrations (42°C, 20 min), ATP was added and then the reactions were shifted to 25°C to initiate refolding. Fluc activity was measured over time and normalized to untreated native Fluc at 0 h. Data are mean ± SD; n = 3 independent replicates. Data were fitted with logistic growth model. Statistical significance relative to the chaperone-containing Raptinal-free condition at the end point was calculated by two-way ANOVA with Dunnett’s correction.

HSP90 promotes refolding of misfolded firefly luciferase and limits its aggregation^27^. We therefore asked whether Raptinal-induced folding stress is buffered by HSP90. Treatment with either Raptinal or the HSP90 inhibitor 17-AAG alone did not induce detectable Fluc-EGFP aggregation, whereas combined treatment induced bright Fluc-EGFP aggregates in approximately 40% of cells (Fig. 2d–f). Misfolded proteins can engage the ubiquitin-proteasome pathway for degradation. Consistently, Raptinal induced a substantial increase in poly-ubiquitinated protein species, which was prevented by CHX (Supplementary Fig. 3c), indicating that this ubiquitin-conjugated burden originates from newly synthesized proteins. Together, these findings indicate that Raptinal impairs the productive folding of newly synthesized proteins, producing misfolded species that are buffered by HSP90-dependent chaperoning and ubiquitin–proteasome–mediated quality control.

If an increased load of misfolded newly synthesized proteins drives mitochondrial damage, then lowering the cell’s capacity to buffer folding stress should sensitize cells to this injury. Indeed, inhibition of cytosolic HSP90 with 17-AAG, cytosolic HSP70 with VER-155008, p97/VCP with CB5083, or the proteasome with MG132, bortezomib, or epoxomicin did not trigger apoptosis within our assay time window, but each significantly potentiated Raptinal-induced apoptosis (Fig. 2g–j). Thus, cytosolic proteostasis pathways buffer this folding stress and limit its progression to mitochondrial apoptosis.

We next asked whether the observed folding defects reflect a direct effect of Raptinal on protein folding. Raptinal consists of two hydrophobic fluorenyl groups connected by a hydrophilic cyclic bis-hemiacetal in aqueous solution (Supplementary Fig. 3d)^18^. Because fluorene derivatives can bind amyloid-β monomers and remodel their aggregation behavior^28^, we hypothesized that Raptinal might directly perturb protein folding. We therefore performed an in vitro refolding assay using recombinant Fluc. Heat denaturation at 42 °C for 10 min abolished luciferase activity (Fig. 2k). When heat-denatured Fluc was allowed to refold at 25 °C in the absence of Raptinal, luciferase activity slowly recovered to approximately 58% of native activity over 24 hours (Fig. 2k). Raptinal slowed refolding and reduced the maximal recovery of luciferase activity in a dose-dependent manner. For example, 1.25 µM Raptinal reduced maximal recovery to approximately 48%, whereas 10 µM Raptinal reduced recovery to approximately 23% (Fig. 2k). In contrast, incubation of native luciferase with up to 20 µM Raptinal for 24 hours did not reduce activity relative to control conditions (Fig. 2l), indicating that Raptinal impairs productive folding without destabilizing the natively folded enzyme — unlike heat, which denatures the mature protein. We further examined whether Raptinal interferes with chaperone-assisted folding by incubating heat-denatured Fluc with a previously established HSP70 chaperone system containing recombinant human HSC70, DNAJB1, and APG2^29^. Denatured Fluc rapidly refolded in the presence of the HSP70 chaperone system, whereas Raptinal strongly impaired this process in a dose-dependent manner (Fig. 2m).

These data indicate that Raptinal directly impairs productive protein folding and thereby induces a proteotoxic state driven by misfolding of newly synthesized proteins.

### Proteotoxic stress is uncoupled from mitochondrial permeabilization by disruption of energy-dependent mitochondrial processes

We next asked how Raptinal-induced proteotoxic stress is coupled to mitochondrial outer membrane permeabilization. Misfolded proteins could compromise mitochondrial integrity from the cytosolic side, for instance, by engaging pore-forming proteins, or they could become deleterious upon entering mitochondria, for example, through energy-dependent protein import. To distinguish these possibilities, we first tested whether permeabilization depends on energy-dependent mitochondrial processes using carbonyl cyanide-p-trifluoromethoxyphenylhydrazone (FCCP), a protonophore that dissipates the mitochondrial membrane potential and broadly perturbs energy-dependent processes, including protein import. In parallel, we examined VBIT4, a small molecule reported to inhibit VDAC oligomerization^30^ and widely used to suppress mtDNA leakage and cGAS–STING signaling in models of aging-associated neuroinflammation and proteostasis-related neurodegeneration^8, 10–12^, although its molecular target and selectivity have recently been questioned^31^.

Unlike FCCP, VBIT4 did not collapse the mitochondrial membrane potential or fragment the mitochondrial network (Supplementary Fig. 4a,b). To exclude a potential contribution of BID or tBID pore-forming activity^32^, we generated pooled BAX/BAK/BID triple knockout HEK293 cells (Supplementary Fig. 4c). In both wild-type and BAX/BAK/BID^TKO^ HEK293 cells, VBIT4 and FCCP prevented Raptinal-induced apoptosis, as assessed by caspase-9 and caspase-3 cleavage (Fig. 3a). This protection was selective for Raptinal, as FCCP and VBIT4 did not suppress BH3-mimetic-induced apoptosis in HeLa cells (Fig. 3b). Additional inhibitors of oxidative phosphorylation, including the complex I inhibitor rotenone, the complex III inhibitor antimycin A, and the ATP synthase inhibitor oligomycin A, also suppressed Raptinal-induced caspase-3/7 activation without affecting the response to ABT-737/S63845 (Supplementary Fig. 4d). Immunofluorescence analysis in CASP9^KO^ HEK293 cells showed that Raptinal-induced SMAC leakage was prevented by VBIT4 and FCCP (Fig. 3c–e), although mitochondrial network fragmentation still occurred (Supplementary Fig. 4e).

**Figure 3.**
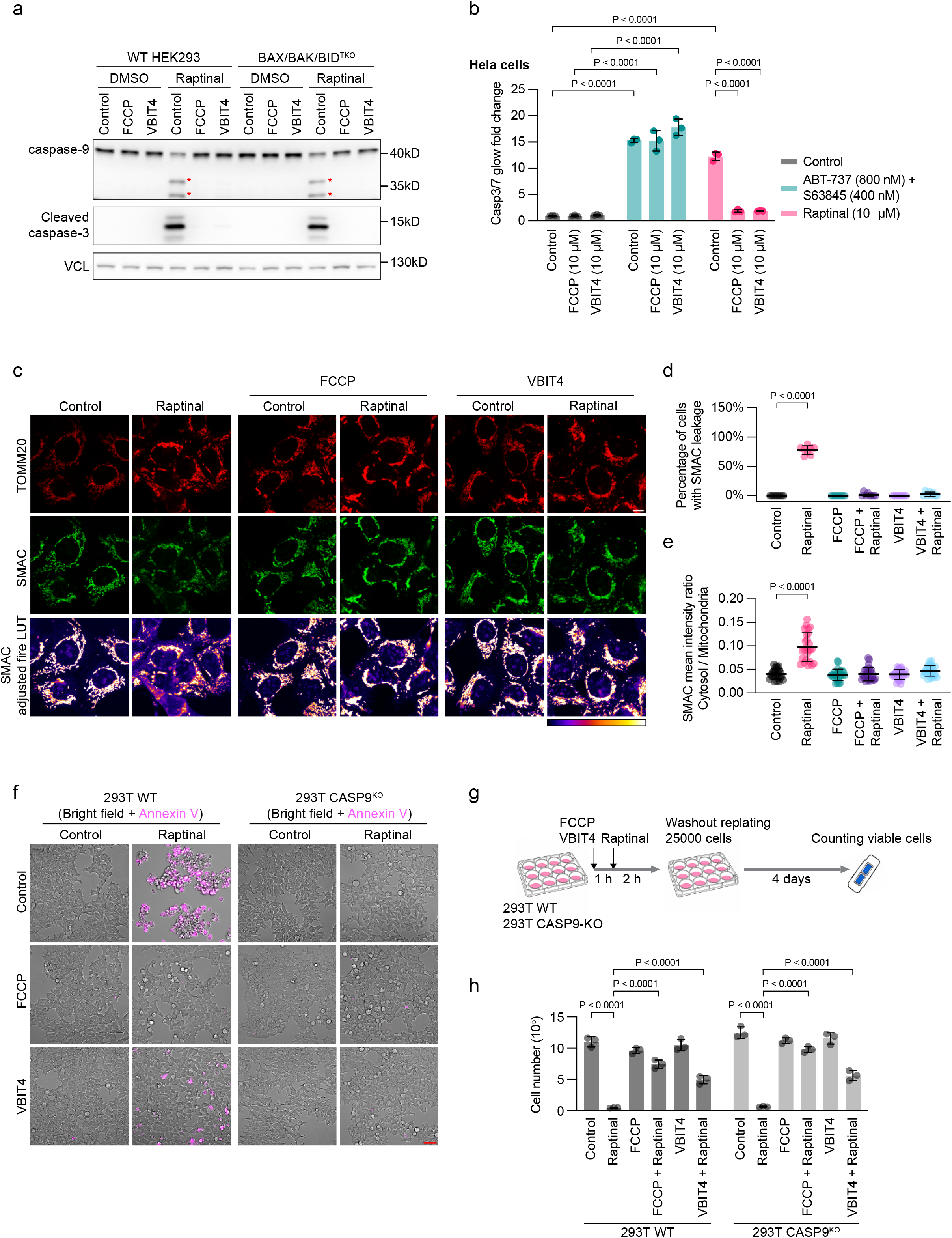
VBIT4 and disruption of oxidative phosphorylation uncouple proteotoxic stress from mitochondrial permeabilization. **(a)** Wild-type and BAX/BAK/BID^TKO^ HEK293 cells were pretreated with vehicle, FCCP (10 µM) or VBIT4 (10 µM) for 1 h and then treated with 10 μM Raptinal for 2 h. Caspase-9 processing and caspase- 3 cleavage were analyzed by western blotting. Cleaved caspase-9 is indicated by red asterisk labels. VCL, loading control. Representative of 2 independent experiments. **(b)** HeLa cells were pretreated with vehicle, FCCP (10 µM) or VBIT4 (10 µM) for 1 h and then treated with Raptinal (10 μM) or BH3 mimetics (ABT-737 at 800 nM and S63845 at 400 nM) for 2 h. Caspase- 3/7 activity was normalized to control condition. Data are mean ± SD; n = 3 independently treated wells, representative of 3 independent experiments; two-way ANOVA with Tukey’s correction. **(c)** CASP9^KO^ HEK293 cells were pretreated with vehicle, FCCP (10 µM) or VBIT4 (10 µM) for 1 h and then treated with DMSO or Raptinal (10 μM) for 2 h. Cells were fixed and immunostained for TOMM20 and SMAC. Representative confocal images are shown with TOMM20 in red and SMAC in green or contrast-enhanced Fire LUT. Scale bar, 10 μm. **(d)** Percentage of cells with increased cytosolic SMAC signal after treatment as in **(c)**. Each point represents one field; bars indicate mean ± SD. n = 8 fields per condition pooled from 3 independent experiments. Statistical significance relative to the control condition was calculated by one-way ANOVA with Dunnett’s correction. **(e)** Quantification of cytosolic-to-mitochondrial SMAC intensity ratio from cells treated as in **(c)**. Each point represents one cell; bars indicate mean ± SD. n = 25 cells per condition pooled from 3 independent experiments. Statistical significance relative to the control condition was calculated by one-way ANOVA with Dunnett’s correction. **(f)** Wild-type and CASP9^KO^ HEK293T cells were pretreated with vehicle, FCCP (10 µM) or VBIT4 (10 µM) for 1 h and then treated with DMSO or Raptinal (10 μM) for 2 h in the presence of Alexa Fluor 647 Annexin V. Representative images are shown as overlay of bright-field and Annexin V fluorescence images. Scale bar, 50 µm. **(g)** Experimental design for **(h)**. Wild-type and CASP9^KO^ HEK293T cells were treated as in **(f)**. After the treatment, 25,000 cells were replated into 12-well plates. Viable cells were counted after 4 days of culture to assess proliferative recovery under indicated conditions. **(h)** Cells were treated as in **(g)**. Viable cell numbers of indicated conditions were counted and plotted as mean ± SD. n = 3 biological replicates. Statistical significance relative to the Raptinal-treated condition for each genotype was calculated by two-way ANOVA with Dunnett’s correction.

Because VBIT4 has been used to suppress mtDNA leakage in settings that do not involve overt apoptosis, we examined whether VBIT4 and FCCP also prevent Raptinal-induced sublethal mitochondrial leakage. Cytosolic mtDNA was quantified from digitonin-released fractions and normalized to total mtDNA in cell pellets^33^. Sublethal Raptinal treatment induced significant mtDNA leakage, which was blocked by both VBIT4 and FCCP (Supplementary Fig. 4f,g). Mitochondrial permeabilization may compromise long-term cell fitness independently of caspase activation^19^. HEK293T cells were highly sensitive to Raptinal (Supplementary Fig. 1c), with most cells undergoing apoptosis within 2 hours, as assessed by morphology and Annexin V labeling (Fig. 3f, left panel). At this time point, CASP9^KO^ cells were largely protected from Raptinal-induced apoptosis (Fig. 3f, right panel). FCCP or VBIT4 largely prevented Annexin V positivity and preserved adherent morphology after Raptinal exposure (Fig. 3f). Following drug washout and replating, Raptinal-treated wild-type and CASP9^KO^ cells failed to proliferate, whereas FCCP- and VBIT4-protected cells showed substantial recovery (Fig. 3g,h). Notably, although VBIT4 suppressed acute apoptotic signaling, it restored only ∼50% of proliferative recovery relative to VBIT4-treated controls (Fig. 3h), suggesting that residual mitochondrial injury not reflected in acute caspase activation nevertheless compromised long-term fitness. These results indicate that mitochondrial permeabilization compromises long-term fitness under proteotoxic stress independently of caspase activation, and that FCCP and VBIT4 preserved mitochondrial integrity sufficiently to support recovery.

Disruption of oxidative phosphorylation has been proposed to suppress intrinsic apoptosis by promoting cristae junction closure^34^. We tested this mechanism in MIC19^KO^ cells, which lack MIC19 and show concomitant depletion of MIC60, two key MICOS components required for cristae junction formation^35^ (Supplementary Fig. 4h). Raptinal still robustly induced caspase-3/7 activation in MIC19^KO^ cells, and this response was blocked by FCCP and VBIT4 (Supplementary Fig. 4i). Thus, the protective effects of FCCP and VBIT4 cannot be explained solely by cristae junction closure. We next asked whether FCCP or VBIT4 suppresses the upstream proteotoxic stress response. FCCP did not significantly alter Raptinal-induced HSPA6 expression, whereas VBIT4 enhanced HSPA6 induction under Raptinal-treated conditions without affecting basal HSPA6 levels (Supplementary Fig. 4j). Together, these data indicate that Raptinal-induced proteotoxic stress can be uncoupled from mitochondrial permeabilization by disrupting oxidative phosphorylation or VBIT4.

### VDAC channels are dispensable for Raptinal-induced apoptosis and VBIT4-mediated protection

To examine the role of VDAC channels in Raptinal-induced mitochondrial permeabilization, we generated single-clone VDAC1/2 double knockout cells and subsequently introduced gRNAs targeting VDAC3 to obtain pooled VDAC1/2/3 triple knockout cells (Fig. 4a). Proteomic profiling confirmed the absence of VDAC1 and VDAC2 and strong depletion of VDAC3 in pooled VDAC^TKO^ cells (Supplementary Fig. 5a,b; Supplementary Table 2). Several additional mitochondrial proteins^36^, including HK1, HK2, BAK1, TMEM70, NME3, and NCBP2AS2 were also reduced by approximately 50% in VDAC^TKO^ cells (Supplementary Fig. 5c). VDAC^TKO^ HEK293 cells remained viable but exhibited reduced proliferation (Supplementary Fig. 5d,e) and markedly dilated mitochondrial morphology (Supplementary Fig. 5f). Unexpectedly, pooled VDAC^TKO^ cells were significantly sensitized to Raptinal-induced apoptosis, which was suppressed by FCCP or VBIT4 (Fig. 4b). To confirm this result, we generated single-clone VDAC^TKO^ cells (Fig. 4c). Consistently, eight independent clonal VDAC^TKO^ cell lines were also sensitized to Raptinal-induced apoptosis and remained responsive to FCCP and VBIT4 (Fig. 4d).

**Figure 4.**
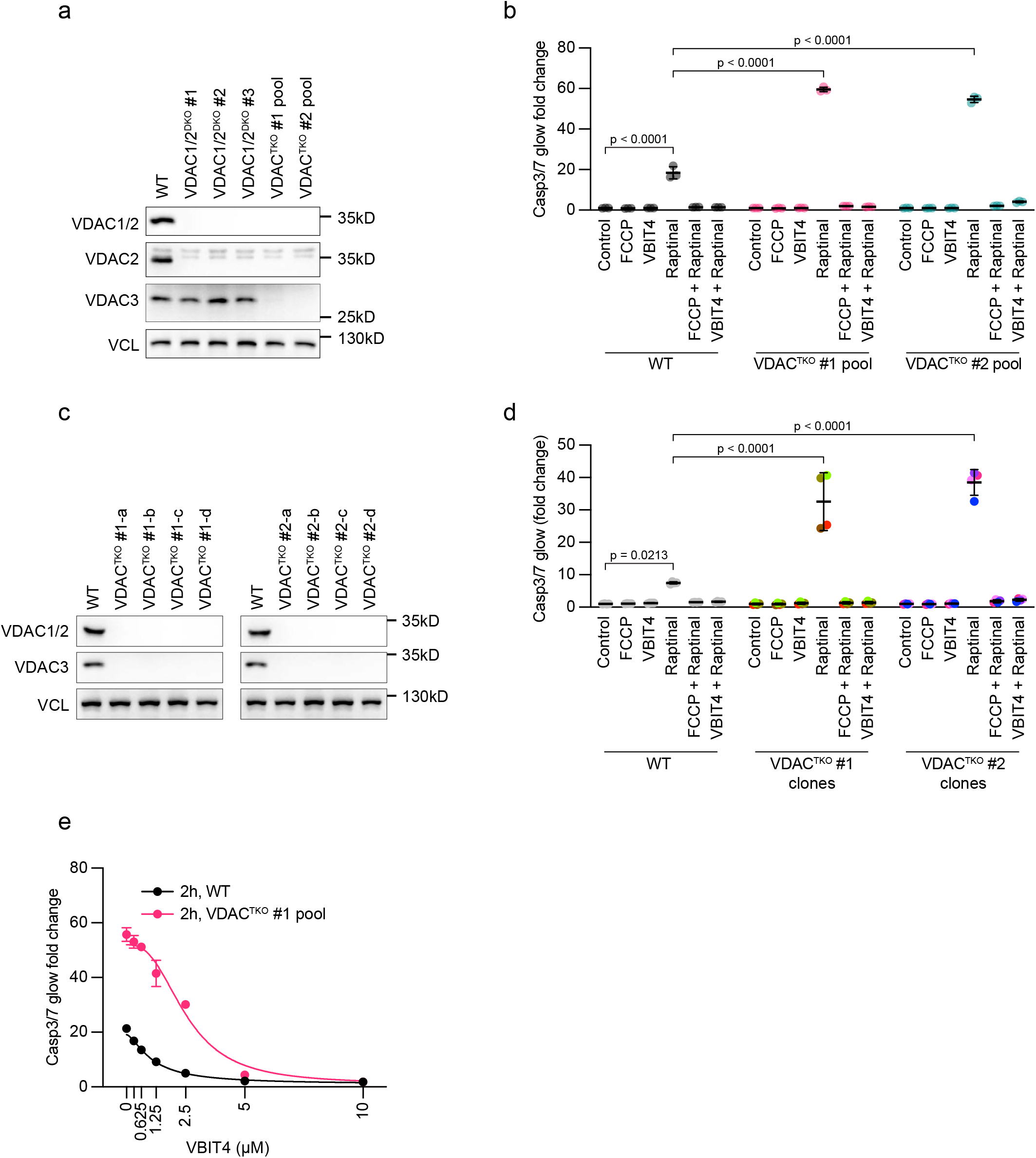
VBIT4 protects mitochondrial integrity independently of VDAC channels. **(a)** Western blot analysis of protein expression of VDAC1, VDAC2 and VDAC3 in wild-type HEK293 cells, three clonal VDAC1/2^DKO^ cell lines (VDAC1/2^DKO^ #1, #2, #3) and two pooled VDAC1/2/3^TKO^ cell lines (VDAC1/2/3^TKO^ #1 pool and #2 pool). VCL, loading control. Representative of 2 independent experiments. **(b)** Wild-type HEK293 cells and two pooled VDAC1/2/3^TKO^ cell lines were pretreated with vehicle, FCCP (10 µM) or VBIT4 (10 µM) for 1 h and then treated with DMSO or Raptinal (10 μM) for 2 h. Caspase-3/7 activity was normalized to control condition of each genotype. Data are mean ± SD; n = 3 independently treated wells, representative of 2 independent experiments; two-way ANOVA with Tukey’s correction. **(c)** Western blot analysis of protein expression of VDAC1/2 and VDAC3 in wild-type HEK293 cells, eight clonal VDAC1/2/3^TKO^ cell lines derived from two pooled VDAC1/2/3^TKO^ cell lines. VCL, loading control. Representative of 2 independent experiments. **(d)** Wild-type HEK293 cells and eight clonal VDAC1/2/3^TKO^ cell lines were pretreated with vehicle, FCCP (10 µM) or VBIT4 (10 µM) for 1 h and followed by DMSO or Raptinal (5 μM) for 2 h. Caspase- 3/7 activity was normalized to control condition of each genotype. Data are mean ± SD; n = 4 for WT cells and 8 independent VDAC1/2/3TKO clonal cell lines, with four clones derived from each parental pool; representative of 2 independent experiments; two-way ANOVA with Tukey’s correction. **(e)** Wild-type HEK293 cells and VDAC1/2/3^TKO^ #1 pool cell lines were pretreated with vehicle or increasing concentrations of VBIT4 for 1 h and then treated with DMSO or Raptinal (10 μM) for 2 h. Caspase-3/7 activity was normalized to control condition of each genotype. Data are mean ± SD; n = 3 independently treated wells, representative of 2 independent experiments. Data were fitted with a four- parameter logistic curve.

In both wild-type and VDAC^TKO^ cells, VBIT4 dose-dependently suppressed Raptinal-induced apoptosis reaching near-maximal protection at 5 µM over the 2-hour assay period (Fig. 4e). At these protective concentrations, VBIT4 did not detectably reduce the mitochondrial membrane potential or induce mitochondrial fragmentation in our assays (Supplementary Fig. 4a,b). Consistently, unlike FCCP or oligomycin A, prolonged VBIT4 treatment did not induce OPA1 processing or OMA1 degradation (Supplementary Fig. 5g), indicating that it did not trigger an inner-membrane stress response^37^. These results indicate that VDAC channels are not required for Raptinal-induced mitochondrial apoptosis or for VBIT4-mediated protection in this system.

We next asked whether VDAC channels are required for VBIT4-dependent suppression of mtDNA leakage. In VDAC^TKO^ cells, cytosolic mtDNA levels were markedly increased (Supplementary Fig. 5h), suggesting that VDAC deficiency predisposes mitochondria to basal mtDNA leakage. Treatment with 5 µM VBIT4 for 24 hours reduced basal mtDNA leakage in wild-type HEK293 cells and significantly decreased cytosolic mtDNA in VDAC^TKO^ cells (Supplementary Fig. 5i). Residual cytosolic mtDNA in VBIT4-treated VDAC^TKO^ cells may reflect incomplete clearance of previously released mtDNA. These data confirm that VBIT4 suppresses mtDNA leakage, as previously reported, and demonstrate that this activity does not require VDAC channels.

### Chemical proteomics links VBIT4 to the TIM23 import machinery

We reasoned that VBIT4 could serve as a chemical probe to identify mitochondrial processes that couple proteotoxic stress to mitochondrial permeabilization. We therefore synthesized ABV22, a photoreactive, clickable VBIT4 analog (Fig. 5a and Supplementary Fig. 6a), which retained VBIT4- like activity in blocking Raptinal-induced apoptosis (Fig. 5b). Cells were labeled with ABV22 at 1 µM for 1 hour. Proximal proteins were covalently captured by UV photocrosslinking, enriched, and identified by quantitative mass spectrometry using label-free intensity-based quantification^38^, with DMSO-treated and non-irradiated (NoUV) samples as controls (Supplementary Table 3). Among the 20 proteins enriched in ABV22-crosslinked samples relative to both controls (log2FC > 3, p < 0.05), 18 were transmembrane proteins (Supplementary Table 4). These data indicate that ABV22 enriches broadly within cellular membranes^31^. Several mitochondrial inner membrane proteins, including TIMM17B, SLC25A20, UQCR10, UQCRQ and FAM162A, were among the most strongly enriched proteins (log2FC > 4 relative to at least one control; Fig. 5c, Supplementary Fig. 6b).

**Figure 5.**
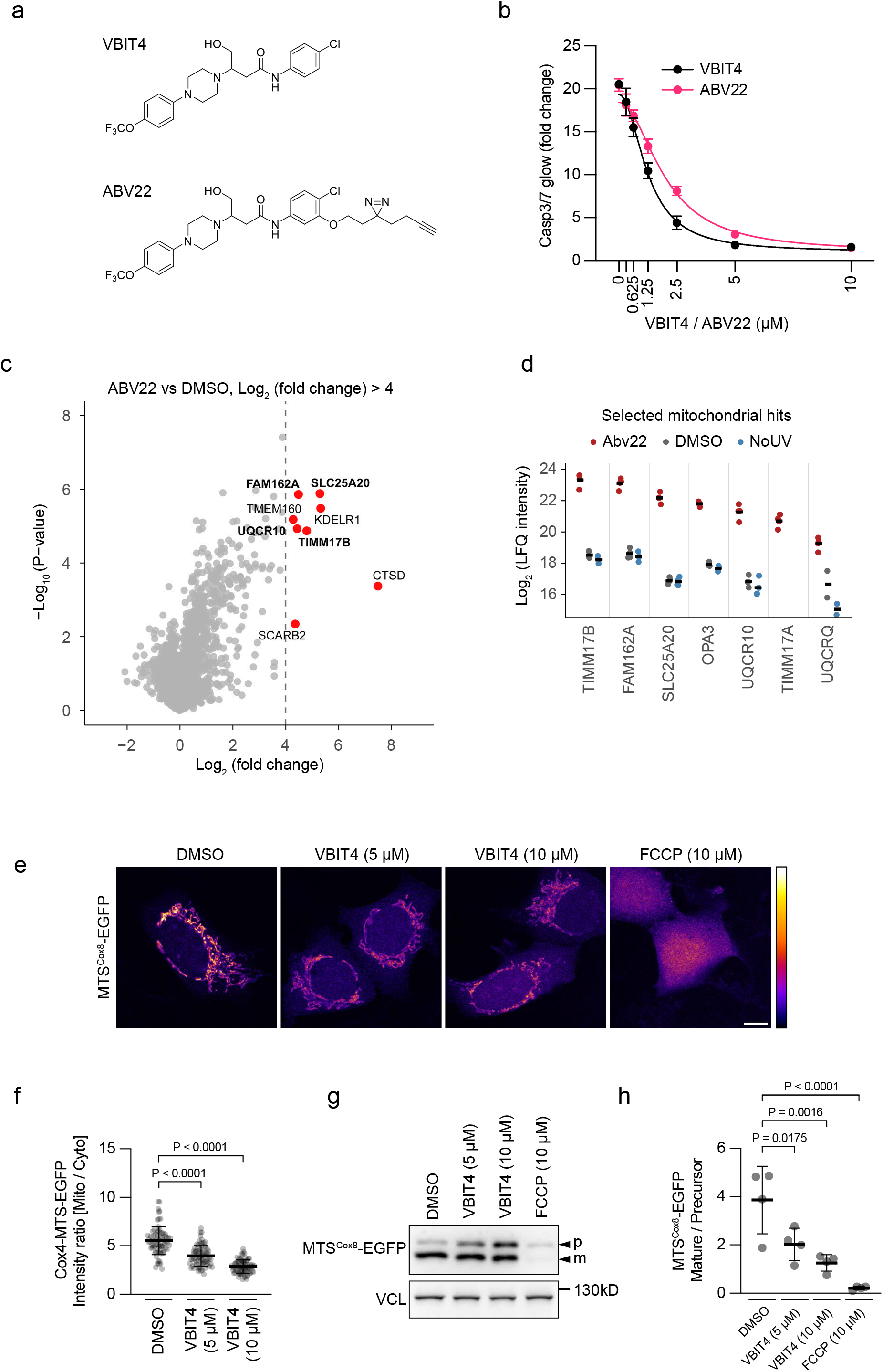
Chemical proteomics links VBIT4 to the TIM23 translocase. **(a)** Chemical structures of VBIT4 and the photoreactive clickable VBIT4 analog ABV22. ABV22 contains a diazirine group for UV-induced photocrosslinking and an alkyne handle for click chemistry. **(b)** Wild-type HEK293 cells were pretreated with vehicle or increasing concentrations of VBIT4 or ABV22 for 1 h and then treated with DMSO or Raptinal (10 μM) for 2 h. Caspase-3/7 activity was normalized to DMSO-treated control. Data are mean ± SD; n = 3 independently treated wells, representative of 2 independent experiments. Data were fitted with a four-parameter logistic curve. **(c)** HEK293 cells were treated with DMSO or ABV22 (1 μM) for 1 h and subjected to UV-induced photocrosslinking, followed by click chemistry, affinity enrichment, and mass spectrometry. Volcano plot shows proteins enriched in ABV22-crosslinked samples compared with DMSO-treated controls. Selected proteins with log2 fold change > 4 are highlighted in red and mitochondrial proteins are labeled in bold. n = 4 biological replicates per condition. p values were calculated from log₂-transformed LFQ intensities using Welch’s t test. **(d)** MS intensities of selected mitochondrial candidate proteins enriched by ABV22 relative to DMSO and NoUV controls. Proteins are ranked according to their MS intensities in ABV22-crosslinked samples. Bars indicate mean values; n = 4 biological replicates per condition. **(e)** Representative confocal images of HEK293 cells transiently expressing MTS^COX8^-EGFP and treated with DMSO, VBIT4 at the indicated concentrations, or FCCP (10 μM) for 6 h. EGFP signal is shown in Fire LUT. Scale bar, 10 µm. **(f)** Quantification of mitochondrial enrichment of MTS^COX8^-EGFP from cells treated as in **(e)**, calculated as the mitochondrial-to-cytosolic EGFP intensity ratio. n = 72 cells per condition pooled from 3 independent experiments; bars indicate mean ± SD. Statistical significance relative to DMSO- treated control was calculated by one-way ANOVA with Dunnett’s correction. **(g)** HEK293 cells transiently expressing MTS^COX8^-EGFP were treated with DMSO, VBIT4 at the indicated concentrations, or FCCP (10 μM) for 6 h. Precursor and mature forms of MTS^COX8^-EGFP were analyzed by western blotting using anti-GFP antibody. VCL, loading control. p, precursor; m, mature. Representative of 4 independent experiments. **(h)** Quantification of the mature-to-precursor MTS^COX8^-EGFP ratio from cells treated as in **(g)**. Each point represents one biological replicate; bars indicate mean ± SD; n = 4 biological replicates. Statistical significance relative to DMSO-treated control was calculated by one-way ANOVA with Dunnett’s correction.

We noted that TIMM17B displayed the highest enrichment among mitochondrial candidates, and TIMM17A was also enriched in ABV22-crosslinked samples but not control samples (Fig. 5d). Consistent with this proximal association, ABV22 preferentially enriched TIMM17A/B relative to components of the TIM22 and TOM complexes, VDAC proteins, and the cristae-junction protein MIC19 (Supplementary Fig. 6c,d). TIMM17A and TIMM17B are core subunits of the TIM23 translocase, which mediates import of most presequence-containing mitochondrial proteins into the matrix and inner membrane in a membrane-potential- and ATP-dependent manner. ^39^. Because precursor proteins remain at least partially unfolded during translocation, this pathway represents a potential vulnerability under folding stress. Moreover, the proximal enrichment of complex III subunit UQCR10 and UQCRQ is consistent with reported effects of VBIT4 on mitochondrial energetics^31, 40^, which would converge on energy-dependent import. These observations prompted us to test whether VBIT4 affects TIM23-mediated protein import.

To assess TIM23-mediated mitochondrial import of presequence-containing proteins, a TIM23 cargo reporter consisting of the mitochondrial targeting signal (MTS) of human COX8 fused to EGFP (MTS^COX8^-EGFP) was transiently expressed in HEK293 cells for 8 h to provide a newly synthesized pool of import substrate. Under control conditions, EGFP predominantly localized to mitochondria, with mean fluorescence intensity more than fivefold higher within mitochondria than in the surrounding cytoplasm (Fig. 5e,f). Treatment with VBIT4 during reporter expression significantly reduced mitochondrial enrichment of EGFP and led to increased cytoplasmic signal (Fig. 5e,f). As expected, FCCP blocked import of MTS^COX8^-EGFP (Fig. 5e). Upon import into the mitochondrial matrix, the COX8 leader peptide is cleaved from EGFP by the mitochondrial processing peptidase. VBIT4 treatment increased levels of the uncleaved precursor in a dose-dependent manner, whereas FCCP abolished presequence cleavage (Fig. 5g,h). As a control, localization of transiently expressed mEmerald-tagged TOMM20 to the mitochondrial outer membrane was not affected by VBIT4 or FCCP (Supplementary Fig. 6e). These data show that VBIT4 partially restricts TIM23-mediated protein import.

### Restricting TIM23 complex-mediated import protects mitochondrial integrity under proteotoxic stress

We next tested whether TIM23 complex-mediated import is involved in mitochondrial injury under Raptinal-induced folding stress. Because the TIM23 complex is essential for cell viability, chronic depletion is not feasible. We therefore established a moderate siRNA-mediated knockdown of TIMM23 in HEK293 and HeLa cells within 48 hours. TIMM23 knockdown was accompanied by decreased levels of TIMM17A and TIMM17B but did not affect SMAC or cytochrome c expression (Fig. 6a). Upon Raptinal treatment, TIMM23-depleted HEK293 cells showed strongly reduced caspase-3/7 activity (Fig. 6b). In HeLa cells, TIMM23 knockdown similarly reduced Raptinal-induced caspase-3/7 activation (Fig. 6c), caspase-3 cleavage (Fig. 6d), and the fraction of cells displaying apoptotic morphology (Fig. 6e), without affecting apoptosis induced by ABT-737/S63845. Moderate TIMM23 knockdown did not significantly alter mitochondrial morphology or membrane potential (Supplementary Fig. 7a,b). To test whether TIMM23 depletion confers protection through activation of the integrated stress response^41^, cells were treated with ISRIB throughout the siRNA experiment. TIMM23 knockdown remained protective in the presence of ISRIB (Supplementary Fig. 7c), indicating that its effect is not mediated by ISR activation.

**Figure 6.**
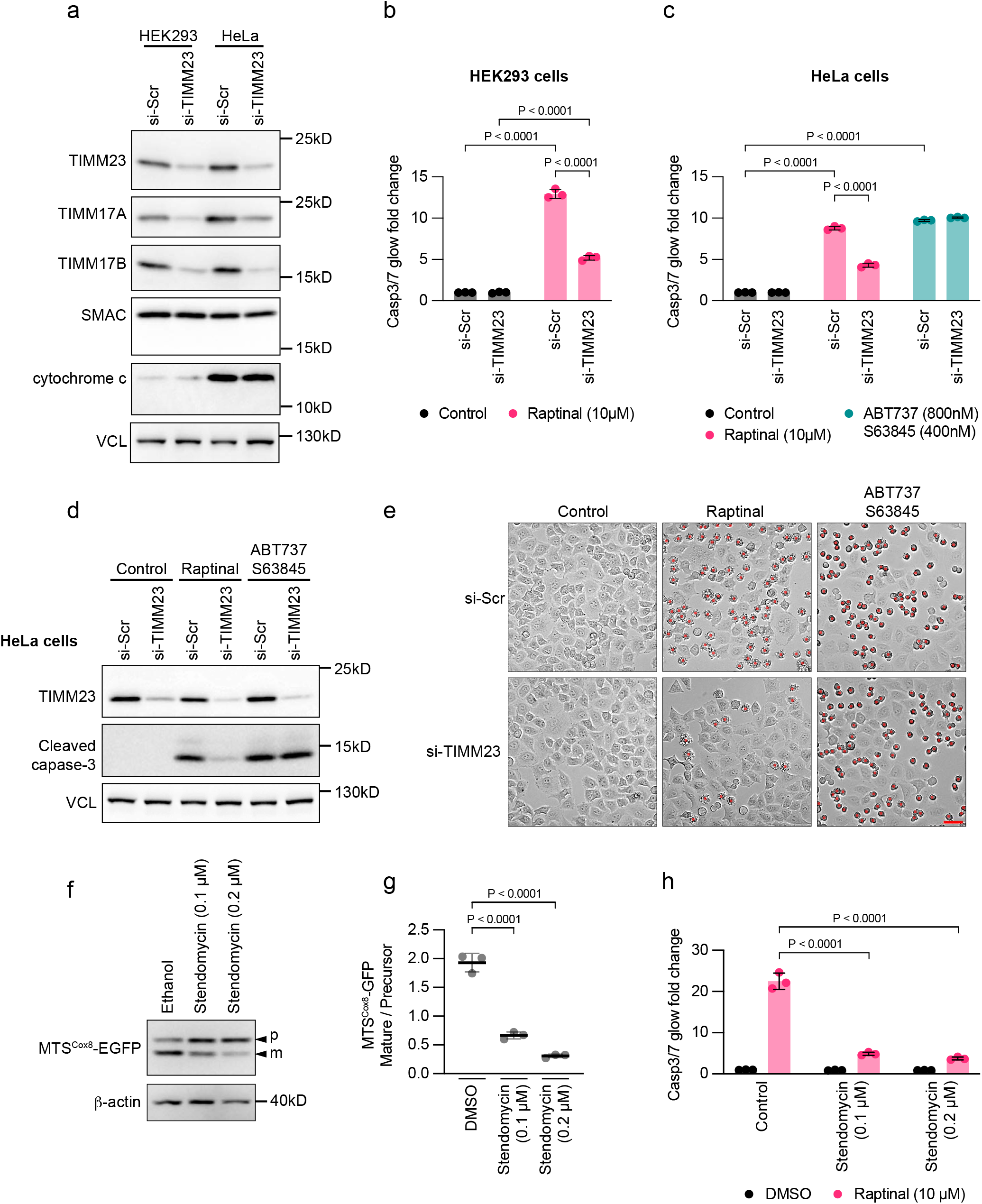
Restricting TIM23 complex-mediated import protects mitochondrial integrity under proteotoxic stress. **(a)** HEK293 and HeLa cells were transfected with non-targeting control siRNA (si-Scr) or TIMM23- targeting siRNA (si-TIMM23) for 48 h. TIMM23, TIMM17A, TIMM17B, SMAC, and cytochrome c expression were analyzed by western blotting. VCL, loading control. Representative of 2 independent experiments. **(b)** HEK293 cells transfected with si-Scr or si-TIMM23 were treated with DMSO or Raptinal (10 μM) for 2 h. Caspase-3/7 activity was normalized to control for either si-Scr or si-TIMM23. Data are mean ± SD. n = 3 independently treated wells, representative of 3 independent experiments; two-way ANOVA with Tukey’s correction. **(c)** HeLa cells transfected with si-Scr or si-TIMM23 were treated with DMSO, Raptinal (10 μM), or BH3 mimetics, ABT-737 (800 nM) and S63845 (400 nM), for 2 h. Caspase-3/7 activity was normalized to control for either si-Scr or si-TIMM23. Data are mean ± SD. n = 3 independently treated wells, representative of 3 independent experiments; two-way ANOVA with Tukey’s correction. **(d)** HeLa cells transfected with si-Scr or si-TIMM23 were treated as in **(c)**. TIMM23 expression and caspase-3 cleavage were analyzed by western blotting. VCL, loading control. Representative of 2 independent experiments. **(e)** Representative bright-field images of HeLa cells transfected with si-Scr or si-TIMM23 and treated as in **(c)**. Cells with apoptotic morphology are manually marked with red dots. Scale bar, 50 µm. **(f)** HEK293 cells transiently expressing MTS^COX8^-EGFP were treated with vehicle or stendomycin at the indicated concentrations for 6 h. Precursor and mature forms of MTS^COX8^-EGFP were analyzed by western blotting using anti-GFP antibody. β-actin was used as loading control. Representative of 3 independent experiments. p, precursor; m, mature. **(g)** Quantification of the mature-to-precursor MTS^COX8^-EGFP ratio from cells treated as in **(f)**. Each point represents one biological replicate; bars indicate mean ± SD. n = 3 biological replicates. Statistical significance relative to control was calculated by one-way ANOVA with Dunnett’s correction. **(h)** HEK293 cells were pretreated with vehicle or stendomycin at indicated concentrations for 5 h and then treated with DMSO or Raptinal (10 μM) for 2 h. Caspase-3/7 activity was normalized to DMSO- treated control. Data are mean ± SD; n = 3 independently treated wells, representative of 3 independent experiments; two-way ANOVA with Tukey’s correction.

To acutely restrict TIM23-mediated import while minimizing adaptive responses to TIMM23 depletion, we used stendomycin, a genetically and biochemically validated inhibitor of the TIM23 complex^42^. At concentrations below 200 nM stendomycin partially restricted TIM23-dependent import of the MTS^COX8^-EGFP reporter (Fig. 6f,g) and suppressed Raptinal-induced apoptosis (Fig. 6h), without affecting mitochondrial morphology or transmembrane potential (Supplementary Fig. 7d,e). Stendomycin also significantly reduced Raptinal-induced SMAC release in CASP9^KO^ cells (Supplementary Fig. 7f,g). At concentrations that partially suppressed Raptinal-induced apoptosis in HeLa cells, stendomycin did not affect BH3-mimetic-induced apoptosis (Supplementary Fig. 7h).

The TIM23 complex is functionally coupled to the presequence translocase-associated motor (PAM), in which HSPA9 facilitates precursor import in an ATP-dependent manner. We therefore asked whether acute pharmacological inhibition of HSPA9 would phenocopy TIM23 restriction. Brief pretreatment with JG-98, a small-molecule HSP70-family inhibitor reported to accumulate in mitochondria and inhibit HSPA9^43^, strongly suppressed Raptinal-induced apoptosis without affecting BH3-mimetic-induced apoptosis (Supplementary Fig. 7i). Taken together, these genetic and pharmacological perturbations establish that restricting the TIM23–PAM import axis protects mitochondrial integrity during translation-associated proteotoxic stress.

## Discussion

Proteostasis decline, mitochondrial dysfunction, and chronic inflammation are recurring features of aging and degenerative disease, yet the mechanistic relationships between these processes remain incompletely understood. Here, we show that impaired folding of newly synthesized proteins compromises mitochondrial integrity independently of canonical pore-forming BCL-2 family proteins but requires mitochondrial protein import. These findings establish a mechanistic link between proteostasis failure and mitochondrial injury and identify protein import as a critical vulnerability during proteotoxic stress.

TIM23-dependent import is essential for mitochondrial proteome maintenance^39^. However, the same pathway imposes a continuous folding and quality-control demand because imported precursors must fold upon arrival. Mitochondrial proteostasis is therefore maintained when the rate of precursor delivery does not exceed matrix folding and quality-control capacity. These two variables are mechanistically coupled: manipulations that lower matrix quality-control capacity, such as inhibition of mitochondrial HSP90 (TRAP1) or depletion of the matrix protease LONP1, reduce TIM23-mediated import^44^, which is expected to restrict precursor influx and limit proteotoxic burden. An acute increase in folding stress, by contrast, may exceed proteostasis capacity on a shorter timescale, exposing a latent vulnerability.

Originally characterized as a rapid inducer of intrinsic apoptosis, Raptinal drives non-canonical mitochondrial permeabilization independently of BCL-2 family pore-forming proteins^18, 19^. Our findings reveal that Raptinal acts as a proteostasis stressor rather than a membrane-permeabilizing agent. It activated integrated stress and heat-shock responses, impaired the folding of newly synthesized proteins, and required ongoing translation to trigger proteotoxic stress and mitochondrial injury. In a luciferase refolding system, Raptinal interfered with both spontaneous and chaperone-assisted refolding without a measurable effect on native luciferase. These data indicate that Raptinal preferentially acts on folding intermediates rather than mature proteins, distinguishing it from heat stress, which also destabilizes mature thermolabile proteins. One possibility is that the fluorene-containing scaffold interacts with folding intermediates that transiently expose hydrophobic surfaces, thereby stabilizing non-productive conformations. Together, these observations establish Raptinal as a distinctive proteostasis stressor that induces acute folding stress on newly synthesized proteins, with mitochondrial permeabilization as one downstream consequence.

The mammalian proteostasis network possesses extensive buffering capacity through chaperone systems, protein-degradation pathways, and adaptive stress responses^15^. The chaperome accounts for up to 10% of total protein mass in human cells^45^, and HSP90 inhibition preferentially affects a subset of sensitive clients^46^. Newly synthesized proteins are particularly susceptible within this hierarchy because they have not yet acquired native conformations while being engaged by subcellular-targeting and quality-control pathways^16^. By impairing productive protein folding at an early stage, Raptinal likely imposes a broad proteotoxic burden that exceeds cellular quality-control capacity, thereby exposing a mitochondrial vulnerability linked to protein import. Consistent with this interpretation, independent perturbations of HSP90, HSP70, VCP/p97, or the proteasome each sensitized cells to Raptinal-induced mitochondrial injury, though none was sufficient on its own to induce comparable injury, likely due to the reserve capacity of the proteostasis network. We therefore favor a model in which mitochondrial protein import constitutes a vulnerability under proteotoxic stress, with Raptinal providing an experimentally tractable trigger.

An entry point into the mechanism linking proteotoxic stress to mitochondrial injury came from VBIT4, originally identified as an inhibitor of VDAC oligomerization and subsequently widely used to suppress mtDNA release and inflammatory signaling in models of aging, neurodegeneration, and sterile inflammation^8, 10–12^. In these settings, VDAC-dependent mtDNA leakage has been proposed as a key pathogenic mechanism. We found that VBIT4 suppressed Raptinal-induced mitochondrial permeabilization and partially restored proliferative recovery, while sparing BH3-mimetic-induced BAX/BAK-dependent apoptosis. However, VBIT4 remained protective in cells lacking all three VDAC isoforms, demonstrating that its protective effects in this context are VDAC independent, consistent with evidence that VBIT4 can act independently of VDAC1^31^. A VBIT4-derived photoaffinity probe enriched TIMM17A/B, core subunits of the TIM23 translocase, and VBIT4 partially restricted TIM23- dependent protein import. Because TIM23-mediated import is energy-dependent, translocase engagement and any energetic effect of VBIT4 would converge on reduced import and cannot be separated in cells. The causal role of import is therefore established independently by orthogonal perturbations of the TIM23/PAM axis: siRNA-mediated depletion of TIMM23, the selective TIM23 inhibitor stendomycin and inhibition of the PAM motor subunit HSPA9 each selectively suppressed Raptinal-induced mitochondrial injury without affecting BH3-mimetic-induced apoptosis. Consistent with the energy dependence of TIM23-mediated import, disruption of oxidative phosphorylation or dissipation of the mitochondrial membrane potential reproduced this protection. This mechanism is distinct from classical mitochondrial import stress, in which impaired translocation and precursor accumulation elicit stress responses. Instead, these findings indicate that the proteotoxic burden becomes deleterious to mitochondria through its coupling to continued TIM23-dependent protein import. Our data caution against inferring VDAC dependence solely from VBIT4 sensitivity and raise the possibility that mtDNA leakage and downstream cGAS–STING signaling in models of aging and neurodegeneration arise, at least in part, from mitochondrial proteostasis defects involving protein import.

The proteotoxic burden imposed on mitochondria can arise through multiple non-mutually exclusive mechanisms. Bona fide mitochondrial precursor proteins are particularly susceptible to folding perturbation since they must remain at least partially unfolded during translocation and acquire their native conformations after import^47, 48^. Alternatively, misfolded non-mitochondrial proteins may expose cryptic targeting information and aberrantly engage mitochondrial import pathways. This possibility conceptually parallels the yeast MAGIC pathway^49^ and related observations in mammalian systems^50^, in which misfolded cytosolic proteins undergo TIM23-dependent mitochondrial import. In either case, mitochondrial injury may ensue when the influx of folding-incompetent polypeptides exceeds mitochondrial proteostasis capacity. Whereas severe disruption of mitochondrial import is expected to compromise mitochondrial homeostasis and induce cytosolic proteotoxic stress, partial attenuation could reduce the proteostasis burden imposed by protein influx, thereby limiting progression to overt mitochondrial permeabilization. Consistent with this idea, delayed mitochondrial protein import protects yeast cells against toxic C-terminal alanine/threonine (CAT)-tailed proteins when mitochondrial ribosome-associated quality control (mitoRQC) fails^51^. Identifying the toxic protein species and their site of action along the import pathway represents the next mechanistic challenge. Our data support a model in which translation-associated proteotoxic stress compromises mitochondrial integrity through the TIM23/PAM-dependent import axis and suggest that regulation of import flux gates whether proteostasis failure propagates to mitochondrial injury.

## Methods

### Cell culture

HEK293, HEK293T, and HeLa cells were maintained in high-glucose DMEM (Gibco) supplemented with 2 mM L-glutamine, 1 mM sodium pyruvate, 10% fetal bovine serum (FBS; Gibco), and 1% penicillin–streptomycin (Gibco). Cells were cultured under sterile conditions at 37°C in a humidified incubator containing 5% CO₂. All cell lines tested negative for mycoplasma contamination.

### Plasmids

For inducible expression of HA-tagged firefly luciferase folding reporters, coding sequence for wild-type firefly luciferase (Fluc-WT) was subcloned from FlucWT-HA-GFP11-N1 (Addgene plasmid #91954) into PB-TRE-dCas9-VPR (Addgene plasmid #63800), replacing the original dCas9-VPR cassette. For inducible expression of EGFP-tagged firefly luciferase folding reporters, the GFP11 fragment in FlucWT-HA-GFP11-N1 was replaced with the EGFP coding sequence, and the resulting Fluc-WT-EGFP cassette was subcloned into PB-TRE-dCas9-VPR, replacing the original dCas9-VPR cassette.

To generate knockout cells by CRISPR-Cas9, sgRNA target sequences were cloned into pSpCas9(BB)- 2A-Puro (PX459) V2.0 (Addgene plasmid #62988) for knockout of CASP8, BAX, and BAK1, or into pMini.puro (generated in-house) for knockout of other genes. The following sgRNA target sequences were used: CASP9, CCGCCGATCCGCTTCGTCCA; MIC19 (gene name CHCHD3), CAGGAACCGGAATCATGGGT; VDAC1, TTTTCTGTTCAGCTTGCACG; VDAC2, TGTTAGGAATTTTCAACGTC; VDAC3, CCATATTTGTACCGAACACA; BID, AGAACCTACGCACCTACGTG; BAX, TCGGAAAAAGACCTCTCGGG; BAK1, GGCGGTAAAAAACGTAGCTG; and CASP8, GCTCTTCCGAATTAATAGAC. All plasmids were verified by Sanger sequencing.

### Antibodies and chemical tools

Antibodies and chemical tools used in this study are listed in Supplementary Table 5.

### Transient transfection and stable cell line generation

siRNA transfection was performed using Lipofectamine 2000 (Invitrogen) according to the manufacturer’s protocol. Briefly, cells were trypsinized, resuspended in Opti-MEM (Gibco), and seeded in 6-well plates at a density of 3 × 10⁵ per well. For each well, 40 pmol ON-TARGETplus Human TIMM23 SMARTpool siRNA (Dharmacon, L-190121-00-0005; target sequences: AAUGAAUGGUCUUCGGCUA, GUUACUCGCACGCGGAUUU, CUGGAGAUCUUGCACGUAU, and CGAUACCUCGUGCAGGAUA) or ON-TARGETplus Non-targeting Control Pool siRNA (Dharmacon, D-001810-10-05) was transfected using Lipofectamine 2000 at the time of seeding. At 6 h after transfection, the medium was replaced with high-glucose DMEM (Gibco) supplemented with 2 mM L-glutamine, 1 mM sodium pyruvate, 10% FBS (Gibco), and 1% penicillin–streptomycin (Gibco). For ISRIB treatment during TIMM23 knockdown, 200 nM ISRIB was added at the time of medium exchange. For western blotting, cell lysates were collected 48 h after transfection. For cell-death assays, transfected cells were replated in 96-well plates at 12,000 cells per well 24 h after transfection and analyzed 48 h after transfection.

Transient plasmid transfection was performed using GeneJuice transfection reagent (Novagen). To generate stable cell lines inducibly expressing firefly luciferase folding reporters, piggyBac plasmids encoding the expression cassette (1 µg) and a plasmid encoding hyperactive piggyBac transposase (0.2 µg) were co-transfected into CASP9^KO^ HEK293 cells in 6-well plates using 2.4 µL GeneJuice reagent. At 24 h after transfection, cells were replated into T25 flasks and selected with 50 µg/mL hygromycin for 1 week.

To generate knockout cells by CRISPR-Cas9, 2 µg PX459 plasmid for knockout of CASP8, BAX, or BAK1, or pMini.puro plasmid expressing the respective sgRNA, was transfected into target cells in 6- well plates using 4 µL GeneJuice reagent per well. At 24 h after transfection, cells were replated into T25 flasks and selected with 2 µg/mL puromycin for 48 h. Surviving cells were either used as knockout pools or single-cell cloned into 96-well flat-bottom plates at a density of 0.8 cells per well. Individual clones were expanded, and knockout efficiency was screened and validated by western blotting.

To generate VDAC1/2^DKO^ cells, plasmids encoding sgRNAs targeting VDAC1 and VDAC2 were mixed at a 1:1 ratio before transfection. Single-cell-derived VDAC1/2^DKO^ clones were screened for loss of puromycin resistance and then transfected with plasmids encoding sgRNA targeting VDAC3 to generate VDAC^TKO^ cells. To generate BAX/BAK^DKO^ cells, plasmids encoding sgRNAs targeting BAX and BAK1 were mixed at a 1:1 ratio before transfection. Pooled BAX/BAK^DKO^ cells were further transfected with plasmid encoding sgRNA targeting BID to generate BAX/BAK/BID^TKO^ cells.

### Mitochondria fractionation and in vitro permeabilization

2 × 10^7^ HEK293 cells were washed once with PBS and resuspended in precooled hypotonic buffer (10 mM KCl, 1.5 mM MgCl₂, 10 mM HEPES-KOH, pH 7.4). After incubation on ice for 2 min to allow cell swelling, the cells were disrupted by 30 strokes using a Dounce homogenizer (Pestle B). The cell homogenate was immediately mixed with 2.5 × mitochondrial isolation buffer (175 mM sucrose, 2.5 mM EGTA, 525 mM mannitol, and 10 mM HEPES, pH 7.4) to restore isotonic conditions. Large cell fragments and debris were removed by two successive centrifugation steps (1300 × g, 5 min, 4 °C). Mitochondria were subsequently pelleted by centrifugation (8,000 × g, 15 min, 4 °C), washed once with mitochondrial isolation buffer (70 mM sucrose, 1 mM EGTA, 210 mM mannitol, and 10 mM HEPES, pH 7.4) and once with mitochondrial resuspension buffer (10 mM HEPES, pH 7.4, 250 mM sucrose). The final mitochondrial pellets were resuspended in mitochondrial resuspension buffer. The yield of isolated mitochondria was quantified using BCA assay.

Isolated mitochondria (10 µg) were incubated with digitonin (0.25%, 0.05%, or 0.025%) or Raptinal (50 µM or 10 µM) at 37 °C for 30 min in a final volume of 50 µL. Following incubation, mitochondria were pelleted by centrifugation (8,000 × g, 15 min, 4 °C). The mitochondrial pellet was resuspended in 50 µL of 2 × Laemmli buffer, while 42 µL of the supernatant was mixed with 8 µL of 6 × Laemmli buffer. Both fractions were briefly boiled before proteins were separated by 16% SDS-PAGE and analyzed by western blotting.

### Caspase-Glo 3/7 assay

Caspase-3/7 activity was quantified using the luciferase-based Caspase-Glo 3/7 Assay System (Promega, #G8090) according to the manufacturer’s instructions. Cells were seeded overnight in Greiner CELLSTAR white 96-well microplates at a density of 12,000 cells per well. Cells were pretreated with the indicated inhibitors for 1 h and then treated with Raptinal or BH3 mimetics (ABT- 737 and S63845) at the indicated concentrations for 2 h to induce cell death. In case of stendomycin, due to the slow target engagement of the lipopeptide, pretreatment was extended to 5 h for HEK293 cells and 24 h for HeLa cells. Cell culture medium alone was used as a blank control. Luminescence readings were background-subtracted using the blank control and normalized to the indicated control conditions to calculate relative fold change of caspase-3/7 activity. Because assay dynamic range varied between independent runs owing to differences in reagent batch and kit age, values were normalized within each experiment and were not pooled across runs. Statistical analysis was performed using independently treated wells on one plate, and results were validated in more than two independent experiments.

### Cell proliferation and proliferative recovery assays

For basal proliferation assays, 25,000 cells were seeded in complete growth medium and cultured under standard conditions. At the indicated time points, cells were detached by trypsinization, resuspended in complete medium, and viable cell numbers were determined by trypan blue exclusion.

For proliferative recovery assays after acute Raptinal treatment, wild-type and CASP9^KO^ HEK293T cells were seeded in 12-well plates and allowed to attach overnight. Cells were pretreated with vehicle, FCCP (10 µM) or VBIT4 (10 µM) for 1 h, followed by treatment with DMSO or Raptinal (10 µM) for 2 h. After treatment, cells were detached by trypsinization, washed twice with PBS and resuspended in drug-free complete medium. For each replicate, 25,000 cells were replated per well in 12-well plates. Cells were then cultured for 4 days under standard conditions to assess proliferative recovery. At the endpoint, cells were detached by trypsinization, and viable cell numbers were determined by trypan blue exclusion.

### Firefly luciferase reporter measurement in cells

CASP9^KO^ HEK293 cells stably expressing doxycycline-inducible HA-tagged wild-type firefly luciferase (Fluc-WT) were seeded overnight in Greiner CELLSTAR white 96-well microplates at a density of 12,000 cells per well. Cells were treated with 200 ng/mL doxycycline for 2 h, followed by addition of Raptinal at the indicated concentrations for an additional 6 h. During Raptinal treatment, doxycycline was maintained at 200 ng/mL. Luciferase activity was measured using the Dual-Glo Luciferase Assay System (Promega, #E2920). Cell culture medium alone was used as a blank control. Luminescence readings were background-subtracted using the blank control and normalized to the indicated control condition.

For analysis of luciferase protein abundance, three parallel wells from each condition were pooled, lysed, and analyzed by western blotting and densitometry. Specific luciferase activity was calculated by dividing normalized luminescence by luciferase protein abundance. To determine whether Raptinal directly interfered with luciferase activity, cells were treated with Raptinal at the indicated concentrations 10 min before addition of the Dual-Glo Luciferase Assay reagent.

### In vitro refolding assay

For spontaneous refolding, recombinant firefly luciferase (Fluc; Promega, #E1701) was diluted to 10 nM in refolding buffer containing 25 mM HEPES/KOH (pH 7.5), 100 mM KCl, 10 mM Mg(OAc)2, 0.05% Tween-20, and 2 mM DTT. Fluc was heat-denatured at 42 °C for 10 min in the presence of the indicated concentrations of Raptinal (0–20 µM). Refolding was initiated by shifting the reactions to 25 °C. To control for effects of Raptinal on native enzyme activity, native Fluc was incubated at 25 °C without heat denaturation in the spontaneous refolding buffer in the presence of the indicated concentrations of Raptinal.

For chaperone-mediated refolding, the recombinant human HSP70 chaperone system has been previously described^29^. Briefly, 10 nM Fluc was heat-denatured at 42 °C for 20 min in buffer containing 25 mM HEPES/KOH (pH 7.6), 100 mM KOAc, 10 mM Mg(OAc)2, 5 mM DTT, 2 µM recombinant human HSC70, 1 µM recombinant human DNAJB1, and 0.2 µM recombinant human APG2, in the presence of the indicated concentrations of Raptinal (0–20 µM) with final DMSO concentration of 0.2%. Refolding was initiated by adding 2 mM ATP and then shifting the reactions to 25 °C to initiate refolding.

At the indicated time points, 1 µL of each refolding reaction was mixed with 24 µL Luciferase Assay System reagent (Promega, #E1500), and Fluc bioluminescence was measured for 2 s using a Lumat LB9508 luminometer (Berthold Technologies). For spontaneous refolding, relative Fluc activity was calculated by normalizing the bioluminescence signal of refolded Fluc to that of native Fluc measured at the corresponding time point. For native Fluc stability measurements and chaperone-mediated refolding, values were normalized to the untreated native Fluc sample at 0 h.

### Immunofluorescence microscopy

For immunofluorescence of fixed cells, cells were cultured on 8-well ibiTreat µ-Slides (ibidi GmbH, #80826) coated with poly-L-lysine (0.01%; Sigma-Aldrich, #P4832). For measurement of mitochondrial transmembrane potential, cells were incubated with 50 nM MitoBrilliant 646 (Tocris, #7700) for 30 min before fixation. Cells were fixed with 4% paraformaldehyde (PFA) for 20 min at room temperature, permeabilized with 0.1% Triton X-100 in PBS for 20 min at room temperature, and blocked with 5% BSA in PBS for 1 h at room temperature. Cells were then incubated overnight at 4 °C with fluorescently conjugated primary antibodies diluted 1:500 in 3% BSA in PBS. Cells were washed three times with PBS to remove unbound antibodies before imaging. Confocal images were acquired at room temperature using a Leica SP8 confocal laser scanning microscope. Image analysis was performed in Fiji.

To evaluate mitochondrial morphology, fixed cells were stained for TOMM20 using either CoraLite594-conjugated TOM20 polyclonal antibody (Proteintech, #CL594-11802) or CoraLite Plus 488-conjugated TOM20 polyclonal antibody (Proteintech, #CL488-11802). Images are shown in grayscale with an inverted LUT.

For quantification of mitochondrial and cytosolic SMAC immunofluorescence signals, cells were stained for SMAC and TOMM20 using CoraLite594-conjugated TOM20 polyclonal antibody (Proteintech, #CL594-11802). Total SMAC signal within each randomly selected cell and SMAC signal within TOMM20-masked mitochondrial regions were quantified in Fiji after background subtraction using cell-free regions. Cytosolic SMAC signal was calculated by subtracting mitochondrial SMAC signal from total cellular SMAC signal, and the cytosolic-to-mitochondrial SMAC ratio was then calculated. To quantify the percentage of cells with SMAC release, random fields were imaged and cells showing elevated cytosolic SMAC signal were counted.

To quantify mitochondrial transmembrane potential, live cells were stained with MitoBrilliant 646 under the indicated conditions. After fixation, cells were stained for SMAC using CoraLite Plus 488- conjugated Smac/DIABLO polyclonal antibody (Proteintech, #CL488-10434) to define the mitochondrial area. MitoBrilliant 646 and SMAC fluorescence intensities were quantified within SMAC-masked mitochondrial regions in randomly selected fields, and the MitoBrilliant 646/SMAC intensity ratio was calculated.

To assess mitochondrial protein import in cells, 4 × 10⁴ HEK293 cells were seeded per well of an 8- well ibiTreat µ-Slide for 16 h. Cells were transfected with 0.05 µg MTS^COX8^-EGFP plasmid per well (Addgene plasmid #172481 mito-mEGFP) using GeneJuice reagent. At 2 h after transfection, cells were treated with VBIT4 or FCCP at the indicated concentrations for an additional 6 h before fixation or collection of cell lysates for western blot analysis. To avoid loss of cytosolic mito-EGFP signal during permeabilization, mitochondria were not immunostained with an additional antibody; instead, mitochondrial regions were masked using the mitochondrial EGFP signal. Fluorescence intensities inside and outside mitochondrial regions were quantified in Fiji. Expression of mEmerald-tagged TOMM20 outer-mitochondrial-membrane protein (mEmerald-TOMM20-N-10; Addgene plasmid #54282) was used as a control.

### Live cell imaging

Live-cell imaging was performed using a 20× objective on a Leica DMi8 live-cell imaging system with environmental control at 37 °C and 5% CO₂. Alexa Fluor 647 Annexin V labeling was used to monitor apoptosis. Wild-type HEK293T or CASP9^KO^ HEK293T cells were seeded in 12-well plates at 1 × 10⁵ cells per well for 16 h. Cells were pretreated with FCCP or VBIT4 at the indicated concentrations for 1 h and then treated with 10 µM Raptinal for an additional 2 h in the presence of Alexa Fluor 647 Annexin V (1:200; BioLegend, #640911).

To assess cell death morphology after TIMM23 knockdown, HeLa cells transfected with either non-targeting control siRNA or TIMM23-targeting siRNA were seeded in 6-well plates at a density of 3 × 10⁵ cells per well. At 48 h after siRNA transfection, cells were treated with Raptinal or BH3 mimetics (ABT-737 and S63845) at the indicated concentrations for 2 h to induce cell death. Cells displaying apoptotic morphology in bright-field images were manually labeled as indicated in the figures.

### Gene expression analysis by RT-qPCR

For gene-expression analysis, RNA was extracted from the indicated cells using TRI Reagent (Sigma-Aldrich, #T9424). RNA samples were diluted to 100 ng/µL, and 500 ng total RNA was used for cDNA synthesis with the iScript cDNA Synthesis Kit (Bio-Rad, #1708890). cDNA corresponding to 5 ng input RNA was used as template for qPCR with Takyon No ROX SYBR 2× MasterMix blue dTTP. qPCR was performed using the following cycling conditions: 95 °C for 3 min, followed by 45 cycles of 95 °C for 30 s, 58 °C for 20 s, and 72 °C for 15 s. All RT-qPCR primers were designed by Integrated DNA Technologies with melting temperatures between 57 °C and 59 °C. All primer sequences are listed in Supplementary Table 5.

### Cytosol extraction and mtDNA quantification

To measure Raptinal-induced mtDNA leakage, CASP9^KO^ HEK293 cells were seeded in a 6-well plate at a density of 2.5 x 10^5^ cells per well. After 24 h, following 1 h pretreatment with 2.5 µM VBIT4 or 10 µM FCCP, cells were treated with 1.25 µM Raptinal for 2 h. For measurement of mtDNA release in VDAC^TKO^ cells, WT and VDAC^TKO^ were treated either with vehicle control or 5 µM VBIT4 for 24 h before cytosolic fraction was extracted.

Cytosolic extraction for mtDNA quantification or western blotting was carried out as previously reported^33^ with minor adaptations. Briefly, cells were collected in PBS and resuspended in 200 µL of digitonin buffer (150 mM NaCl, 20 mM HEPES, 0.002% digitonin) supplemented with cOmplete Mini EDTA-free (Roche, #04693159001) for 20 min at 4 °C under constant rotation. Cell suspension was centrifuged at 4 °C for 5 min at 1,000 × g. Cytosolic extract was collected from the supernatant and was further cleared by centrifugation for 10 min at 17,000 × g. Clarified cytosolic extract was directly used for qPCR analysis or mixed with Laemmli buffer for SDS-PAGE and western blotting. The pellets containing the total mtDNA fraction were resuspended in 500 µL of 50 mM NaOH and boiled for 30 min at 95 °C. The pH was then neutralized with 100 µL Tris-HCl pH 8 and total mtDNA fraction was diluted 1:400 before use. Quantitative PCR was performed on both cytosolic and pellet fractions using validated primers for Dloop and ND1 regions. mtDNA in the cytosolic fraction was normalized to the mtDNA in the pellet fraction for each condition.

### SDS-PAGE and western blot analysis

For protein analysis by western blotting, cells were lysed in RIPA buffer containing 50 mM Tris-HCl, pH 7.4 at 4 °C, 150 mM NaCl, 1% NP-40, 0.5% sodium deoxycholate, and 0.2% sodium dodecyl sulfate, supplemented with cOmplete Mini EDTA-free protease inhibitor cocktail (Roche, #04693159001). Lysates were briefly sonicated and cleared by centrifugation at 15,000 × g for 20 min at 4 °C. Protein concentrations in the supernatants were determined using a BCA assay. Protein concentrations were normalized across samples, and lysates were mixed with 4× Laemmli buffer supplemented with 10 mM TCEP.

For SDS-PAGE, 5–20 µg protein was loaded per sample. All SDS-PAGE was performed under reducing conditions, followed by transfer to PVDF membranes using a wet-tank transfer protocol. PVDF membranes were blocked with 5% BSA or 5% milk in PBST (0.05% Tween-20) and incubated with primary antibodies overnight at 4 °C. Unless otherwise indicated, primary antibodies were used at a dilution of 1:2,000. Western blots were either run in parallel or probed at different regions of the same PVDF membrane. Membrane stripping and antibody re-probing were not performed in this study. Densitometric analysis was performed using Bio-Rad Image Lab software (version 6.0).

### RNA-seq analysis

CASP9^KO^ HEK293 cells were treated with DMSO or Raptinal (10 μM) for 6 h, with three biological replicates per condition. At the end of treatment, cells were lysed directly in TRI Reagent (Sigma-Aldrich, #T9424). RNA was isolated from the TRI Reagent homogenate using the Maxwell RSC48 instrument and the Maxwell RSC miRNA Tissue Kit (Promega). Total RNA purity was assessed using a NanoDrop spectrophotometer, and RNA integrity was analyzed using a Bioanalyzer (Agilent). cDNA libraries were prepared from 100 ng total RNA using the CORALL mRNA kit (Lexogen) according to the manufacturer’s protocol. Individual libraries were barcoded by PCR with unique dual-index primers (Lexogen), pooled in equimolar amounts, and subjected to 60-bp paired-end sequencing on a NextSeq 2000 instrument (Illumina).

RNA-seq data analysis was carried out on the Galaxy server^52^. Paired-end RNA-seq reads were adapter- and quality-trimmed using Cutadapt v5.2^53^ in Galaxy. The 3′ adapter sequence AGATCGGAAGAG was removed from both read 1 and read 2 using a maximum error rate of 0.1 and a minimum overlap of 3 nt. Low-quality bases were trimmed using a Q20 cutoff. Reads shorter than 20 nt after trimming were discarded. Quality check was performed using MultiQC v1.33^54^. Cutadapt-trimmed paired-end reads were aligned to the human hg38 reference genome using RNA STAR v2.7.11b^55^ in Galaxy with default alignment parameters. The built-in hg38 index was used without additional GTF annotation. Alignments were exported as BAM files. Gene-level read counts were generated from STAR-aligned BAM files using featureCounts v2.1.1^56^ in Galaxy with the built-in hg38 annotation. Reads were counted as unstranded paired-end fragments, requiring both reads of a pair to be aligned. The resulting gene-count table was used for differential expression analysis. Differential expression analysis was performed from raw gene counts using the DESeq2^57^ Galaxy tool (2.11.40.8+galaxy2), with treatment as the primary factor. Genes with fewer than 10 counts were pre-filtered before analysis. DESeq2- normalized counts were exported for visualization and downstream summary analyses, while variance-stabilizing transformed counts were used to generate gene-wise z-score heatmaps. Functional enrichment analysis was performed using g:Profiler^58^ in Galaxy (0.1.7+galaxy11) for *Homo sapiens*. Significantly enriched terms were identified using Benjamini–Hochberg FDR correction with a threshold of 0.05. Reactome pathway terms were used for pathway enrichment analysis. For visualization and reporting, broad biological processes with term size >200 and very small signaling branches with term size <10 were excluded.

### Chemical proteomics of ABV22

#### ABV22 labeling, cell lysis, and click chemistry

All proteomics experiments were performed with four biological replicates per condition. Before cell seeding, 6-cm dishes were coated with 0.1 mg/mL poly-L-lysine solution (0.01%, w/v) for 10 min at room temperature, washed with PBS, and dried. HEK293 cells were seeded at 1.5 × 10⁶ cells per 6-cm dish and grown to approximately 90% confluence.

Cells were washed with PBS and then labeled for 1 h under standard culture conditions with ABV22 (1 µM; 1:200 dilution from a DMSO stock) or 0.5% DMSO. After labeling, medium was removed and cells were washed twice with precooled PBS. Cells were irradiated on cooling packs at 365 nm for 5 min. For NoUV controls, ABV22-labeled cells were processed in parallel without UV irradiation.

After irradiation, PBS was removed and cells were lysed directly on the dish with 200 µL lysis buffer containing PBS, pH 7.4, 1% Triton X-100, and 0.5% SDS, supplemented with cOmplete Mini EDTA-free protease inhibitor cocktail (Roche, #04693159001). Cells were lysed for 15 min on ice and transferred to 1.5-mL Eppendorf tubes. Lysates were sonicated twice for 10 s at 20% intensity using a Sonopuls HD 2070 ultrasonic homogenizer (Bandelin electronic GmbH) and cleared by centrifugation at 21,000 × g for 15 min at room temperature. Cleared lysates were transferred to fresh 1.5-mL Eppendorf tubes, and protein concentration was determined by BCA assay (Rotiquant; Carl Roth).

For each sample, protein concentration was adjusted to 2.23 mg/mL in a total volume of 45 µL in a polypropylene V-bottom 96-well plate (Greiner, #651201). Click chemistry was initiated by adding 4.9 µL click-mix per sample, consisting of 0.6 µL 20 mM biotin-azide in DMSO, 2.5 µL 1.67 mM tris(benzyltriazolylmethyl)amine (TBTA) in 80% tert-butanol and 20% DMSO, 1.2 µL 50 mM CuSO₄ in H₂O, and 0.6 µL 100 mM tris(2-carboxyethyl)phosphine (TCEP) in H₂O. Reactions were incubated for 1.5 h at room temperature with shaking at 950 rpm and quenched by adding 65 µL 8 M urea supplemented with 10 mM TCEP and 20 mM iodoacetamide (IAA) per sample. After incubation for 15 min at 25 °C with shaking at 950 rpm, residual IAA was quenched by adding 2 µL 500 mM dithiothreitol (DTT) in H₂O per sample.

#### Enrichment, digestion, and peptide cleanup

All reagents used for sample processing were LC-MS grade. Sample processing was adapted from Mostert et al.^59^. For each sample, 10 µL of a 2× concentrated 1:1 mixture of washed hydrophobic and hydrophilic carboxylate-coated magnetic beads (Cytiva; washed three times with H₂O) was added. Proteins were precipitated onto the beads by adding 175 µL ethanol per sample. The plate was then transferred to an automated liquid-handling system (Hamilton Microlab Prep) for further processing and incubated for 5 min with shaking at 500 rpm.

For each wash step, the plate was placed on a 96-well ring magnet (Alpaqua Magnum FLX) for 90 s, and the supernatant was removed at a low aspiration speed of 20 µL/s to avoid bead loss. The plate was then removed from the magnet, the next wash solution was added, and the samples were shaken for 1 min at 800 rpm. Samples were washed three times with 180 µL 80% ethanol and once with 180 µL acetonitrile.

To elute proteins from the carboxylate-coated beads, 75 µL 0.2% SDS in PBS was added to each sample, and the plate was incubated for 5 min at 40 °C with shaking at 800 rpm. The plate was placed on the magnet, and the supernatant was transferred to new wells. This elution step was repeated once, resulting in a final eluted protein volume of 150 µL per sample.

Meanwhile, streptavidin magnetic beads (New England Biolabs, #S1420S) were washed three times with 0.2% SDS in PBS. Washed streptavidin beads were added to each eluted protein sample at 50 µL per well. The plate was removed from the liquid-handling system, sealed, and incubated for 1 h at 25°C with shaking at 950 rpm in a plate shaker equipped with a heated lid to prevent condensation, allowing biotinylated proteins to bind to the streptavidin beads.

After binding, the plate seal was removed and the plate was returned to the liquid-handling system. Beads were washed three times with 180 µL 0.1% NP-40 in PBS, twice with 180 µL 6 M urea, and three times with 200 µL H₂O. Bead-bound proteins were digested overnight at 37 °C in 100 µL 50 mM triethylammonium bicarbonate (TEAB) containing 0.75 µg sequencing-grade trypsin (Promega), using a tightly sealed plate with a heated lid and shaking at 950 rpm.

After tryptic digestion, peptides were eluted from the beads using the liquid-handling system and desalted on StageTips containing two layers of SDP-RPS material (Empore, 3M). Desalting was performed as previously described by Coscia et al.^60^, with minor modifications. Briefly, StageTips were equilibrated with 150 µL wash buffer 1 consisting of 1% trifluoroacetic acid (TFA) in isopropanol before sample loading. Samples were loaded by centrifugation at 500 × g for 10 min, followed by washing with 170 µL wash buffer 1 at 800 × g for 10 min and 170 µL wash buffer 2 consisting of 0.2% TFA in H₂O. Peptides were eluted with 50 µL elution buffer consisting of 1% ammonia and 80% acetonitrile by centrifugation at 300 × g for 5 min, followed by 800 × g for 5 min. Samples were dried in a centrifugal evaporator and reconstituted in 50 µL 1% formic acid for LC-MS analysis on an Orbitrap Eclipse Tribrid instrument (Thermo Fisher Scientific) operated in data-independent acquisition mode.

#### LC-MS/MS measurements

Peptide quantification was performed using an HPLC-MS/MS system consisting of a Vanquish Neo UHPLC system coupled to an Orbitrap Eclipse Tribrid mass spectrometer (Thermo Fisher Scientific). The Vanquish Neo UHPLC was equipped with a PepMap Neo 5 µm C18 trap cartridge, 300 µm × 5 mm (Thermo Fisher Scientific), and operated in trap-and-elute injection mode, in which samples were loaded onto the trap cartridge before chromatographic separation.

The system was operated at a flow rate of 400 nL/min using buffer A, consisting of 0.1% formic acid (FA) in water, and buffer B, consisting of 0.1% FA in acetonitrile. Peptides were separated on an Aurora Ultimate separation column, 3rd generation, 25 cm, nanoflow UHPLC-compatible (IonOpticks), maintained at 40 °C and coupled to a Nanospray Flex Ion Source (Thermo Fisher Scientific). The HPLC method comprised a 45-min gradient, starting with an increase from 5% to 22% buffer B over 30 min, followed by an increase to 32% buffer B over 5 min and an isocratic wash at 90% buffer B for 10 min. Separation-column washing and equilibration were performed with fast equilibration enabled, using an equilibration factor of 3 at 5% buffer B. Trap-column washing and equilibration were performed with fast wash and equilibration enabled, together with zebra wash using two wash cycles and an automatic equilibration factor.

The Orbitrap Eclipse mass spectrometer was operated in data-independent acquisition mode with internal real-time mass calibration using a user-defined positive lock mass of m/z 445.12003. Full MS scans were acquired in the Orbitrap at a resolution of 60,000, with an AGC target of 4 × 10⁵, a maximum injection time of 100 ms, and a scan range of m/z 400–1,000. MS2 spectra were acquired in the Orbitrap at a resolution of 15,000, with an AGC target of 1 × 10⁶ and a maximum injection time of 40 ms. Quadrupole isolation was performed using 10 m/z windows with 1 m/z overlap across a scan range of m/z 145–1,450. Fragmentation was performed by higher-energy collision-induced dissociation (HCD) using a normalized collision energy of 30%, and fragment ions were detected in the Orbitrap. Data acquisition was performed using Thermo Scientific Foundation software version 3.1 SP9 and Xcalibur version 4.6.

#### Proteomics data analysis

Acquired raw files were converted to mzML format using the MSConvert tool, version 3.0.25054- 207c92d, from ProteoWizard software, version 3.0.21193 64-bit. MS data were processed using DIA-NN, version 1.8.1, in library-free mode^61^. The UniProt reference proteome for *Homo sapiens* (taxon ID: 9606; downloaded on September 9, 2024) was used for library generation.

For precursor ion generation, FASTA digest was enabled for library-free search and library creation, and deep-learning algorithms were used to predict spectra, retention times (RTs), and ion mobilities (IMs). Trypsin/P was specified as the protease, with a maximum of two missed cleavages. N-terminal methionine excision was enabled, carbamidomethylation of cysteines was specified as a fixed modification, and no variable modifications were allowed. Peptide length was restricted to 7–30 amino acids, and precursor charge states were restricted to 2–4. The precursor m/z range was set to 300–1,800, and the fragment m/z range was set to 200–1,800. The precursor false-discovery rate (FDR) was set to 0.01.

Mass accuracy, MS1 accuracy, and scan-window settings were all set to 0. Isotopologues, match-between-runs (MBR), and removal of likely interferences were enabled. The neural-network classifier was operated in single-pass mode. Protein inference was performed at the gene level with heuristic protein inference enabled (--relaxed-prot-inf). Quantification was performed using the robust LC high-precision strategy. Cross-run normalization was RT-dependent, smart profiling was used for library generation, and optimal settings were used for both speed and RAM usage.

#### Statistical analysis of proteomics data

Statistical analyses were performed on log₂-transformed LFQ intensities without imputation of missing values. Statistical significance was assessed on a per-protein basis using a two-sample Welch’s t test. Proteins were tested only when at least two valid values were available in each group. For volcano plot visualization, the y axis was defined as −log₁₀(p value) using the nominal p value. Proteins meeting both criteria, log₂ fold change > 4 and unadjusted p < 0.05, were highlighted as hits and colored according to mean ABV22 log₂ intensity.

### ABV22 labeling in HEK293 cells for western blotting

The experiment was performed with four biological replicates. Before cell seeding, 6-cm dishes were coated with 0.1 mg/mL poly-L-lysine solution (0.01%, w/v) for 10 min at room temperature, washed with PBS, and dried before use. HEK293 cells were seeded at 1.5 × 10⁶ cells per 6-cm dish and grown to approximately 90% confluence. Cells were washed with PBS and incubated for 1 h at 37 °C and 5% CO₂ with either 1 µM ABV22 in 1.5 mL serum-free medium or a medium control. After incubation, medium was removed and cells were washed twice with pre-cooled PBS. Probe-treated dishes were irradiated on cool packs at 365 nm for 5 min.

After irradiation, PBS was removed and cells were lysed directly on the dishes by adding 200 µL lysis buffer containing PBS, pH 7.4, 1% Triton X-100, and 0.5% SDS, supplemented with one tablet of cOmplete Mini EDTA-free protease inhibitor cocktail (Roche, #04693159001) per 10 mL lysis buffer. Cells were lysed for 15 min on ice and transferred to 1.5-mL Eppendorf tubes. Lysates were sonicated once for 10 s at 20% intensity using a Sonopuls HD 2070 ultrasonic homogenizer (Bandelin electronic GmbH) and cleared by centrifugation at 21,000 × g for 15 min at room temperature. Cleared lysates were transferred to fresh 1.5-mL Eppendorf tubes, and protein concentration was determined by BCA assay (Rotiquant; Carl Roth).

For each sample, protein concentration was adjusted to 2.23 mg/mL in a total volume of 90 µL in 1.5- mL Eppendorf tubes. Click chemistry was initiated by adding 9.8 µL click-mix per sample, consisting of 1.2 µL 20 mM biotin-azide in DMSO, 5 µL 1.67 mM TBTA in 80% tert-butanol and 20% DMSO, 2.5 µL 50 mM CuSO₄ in H₂O, and 1.2 µL 100 mM TCEP in H₂O. Reactions were incubated for 1 h at room temperature with shaking at 950 rpm and quenched by adding 600 µL ice-cold acetone. Samples were frozen overnight at −20 °C to precipitate proteins. Proteins were then pelleted by centrifugation at 21,000 × g for 20 min at 4 °C, and the supernatant was removed. Samples were dried for 10 min to remove residual acetone and reconstituted in 200 µL buffer containing 50 mM Tris-HCl, pH 8.0, 150 mM NaCl, and 1% SDS by heating to 90 °C for 10 min with shaking at 300 rpm. Samples were stored at −20 °C.

For affinity purification of biotinylated proteins, 200-µL samples were thawed, boiled at 90 °C for 30 min with shaking at 300 rpm, and centrifuged at 8,000 rpm for 10 min at room temperature. The soluble fraction was diluted in PBS to reduce the final SDS concentration to 0.1% and incubated with Dynabeads MyOne Streptavidin T1 (Invitrogen, #65601) for 3 h at room temperature with constant rotation. The unbound fraction was removed, and beads were washed three times with 1 mL washing buffer containing 50 mM Tris-HCl, pH 8.0, 150 mM NaCl, and 0.05% SDS. Bound proteins were eluted by boiling the beads in 50 µL 1× Laemmli buffer at 90 °C for 10 min.

### Whole cell proteomic profiling

For whole-cell proteomic profiling, 1 × 10⁶ cells per replicate were lysed in 0.3 mL lysis buffer containing 4% deoxycholate and 100 mM Tris-HCl, pH 8.5. Protein concentrations were determined using the Pierce BCA Protein Assay Kit (Thermo Fisher Scientific). Aliquots containing 10 µg total protein were adjusted to a final volume of 5 µL in 100 mM Tris-HCl, pH 8.5, reduced with dithioerythritol at a final concentration of 50 mM for 30 min at 37 °C, and subsequently carbamidomethylated with iodoacetamide at a final concentration of 100 mM for 30 min at room temperature. Proteins were first digested with 100 ng Lys-C (Fujifilm Wako) for 4 h at 37 °C, followed by overnight digestion with 200 ng sequencing-grade modified trypsin (Promega). Digestion was stopped by addition of formic acid to a final concentration of 1%. Before LC-MS/MS analysis, peptides were purified using SCX StageTips (Thermo Fisher Scientific) according to the manufacturer’s recommended protocol.

LC-MS/MS analysis was performed using a nanoElute 2 chromatography system coupled to a timsTOF HT mass spectrometer (both Bruker Daltonics). Peptide samples were loaded in solvent A, consisting of 0.1% formic acid in water, onto a trap column, PepMap Neo Trap Cartridge, 300 µm × 5 mm, 5 µm particles (Thermo Fisher Scientific), and separated on an Aurora Ultimate C18 UHPLC column, 25 cm × 75 µm ID (IonOpticks), at a flow rate of 250 nL/min. Peptides were separated using the following gradient: 2%–25% solvent B, consisting of 0.1% formic acid in acetonitrile, over 25 min; 25%–37% solvent B over 5 min; 37%–95% solvent B over 1 min; followed by equilibration at 95% solvent B for 4 min. MS acquisition was performed using a dia-PASEF method with the following settings: mass range, m/z 475–1,000; ion-mobility range, 0.85–1.27 1/K₀; eight MS/MS ramps; 21 MS/MS windows; and an estimated cycle time of 0.95 s. The mass spectrometry data were deposited in PRIDE.

Proteomics data were analyzed in R. For global comparison of wild-type and VDAC^TKO^ cells, pairwise comparisons of each protein were performed between wild-type cells and each of the two pooled VDAC^TKO^ cell lines. Within each comparison, missing values were excluded on a per-condition basis, and proteins with fewer than two quantitative values in an entire condition were assigned missing test results. Statistical significance was assessed independently for each protein using an unpaired two-sample Student’s t test on LFQ intensities. No multiple-testing correction or value imputation was applied. Log₂ fold changes were visualized in a two-dimensional scatter plot. Proteins with nominal p value < 0.05 and log₂ fold changes > 0.8 or < −0.8 were highlighted in red. Selected mitochondrial proteins shown in Supplementary Fig. 5c were analyzed separately by two-way ANOVA with Dunnett’s correction for multiple comparisons, as indicated in the figure legend.

### Statistics and reproducibility

Statistical analyses were performed using GraphPad Prism version 9.00 (GraphPad Software) and R version 4.5.2 in Positron version 2026.05.1 build 2 (Posit Software). Statistical methods used to predetermine sample size were not performed. Datasets were analyzed using Student’s t tests, one-way ANOVA, or two-way ANOVA, with corrections for multiple comparisons as indicated in the figure legends. Data are presented as mean ± SD, as indicated in the figure legends. Only p values for relevant comparisons are shown.

## Data availability

RNA-sequencing data generated in this study have been deposited in the Gene Expression Omnibus (GEO) under accession number GSE336837. The proteomic dataset of the VDAC^TKO^ cell line has been deposited in the MassIVE repository under accession number MSV000102272 and in the ProteomeXchange Consortium under accession number PXD080174. The proteomic dataset from the ABV22 crosslinking experiments has been deposited in the ProteomeXchange Consortium under accession number PXD080175. Additional information required to reanalyze the data reported in this study is available from the corresponding author upon reasonable request.

## Supporting information

Supplementary Table 5 | Antibody, chemical tool and oligonucleotides

Supplementary Table 4 | ABV22 high enrichment hits

Supplementary Table 3 | ABV22 chemical proteomics

Supplementary Table 2 | VDAC-TKO MS raw intensity file

Supplementary Table 1 | DESeq2 analysis of Raptinal-treated cells

## Acknowledgements

We thank Dr. Thomas Fröhlich and his team in Laboratory for Functional Genome Analysis (LAFUGA, LMU) for MS proteomic analysis. We thank Dr. Dominic Hoepfner (Novartis, Basel, Switzerland) for providing Stendomycin. We thank Maximilian Schuh for his help with the PRIDE upload. We thank Dr. Stefan Krebs and his team in LAFUGA for RNA-seq analysis.

## Author contributions

Z.S. and V.H. conceptualized the study, curated and analyzed data, and wrote the manuscript. Z.S. performed most experiments. H.H. carried out the in vitro refolding experiments. S.B. and A.B. synthesized the chemical probe and performed chemical-proteomics analysis. S.J.S. contributed in vitro mitochondrial assays and cell-death analyses. G.G. contributed western blotting, RT-qPCR analysis of gene expression, and mtDNA analysis. G.S. contributed mtDNA quantification. B.W. provided Stendomycin and protocol for its usage. S.A.S. supervised the chemical-proteomics workflow and data analysis. F.U.H. supervised the in vitro protein-refolding experiments and data analysis.

