## Supplementary Figures 1-7 for "Mitochondrial protein import couples proteostasis failure to mitochondrial permeabilization"

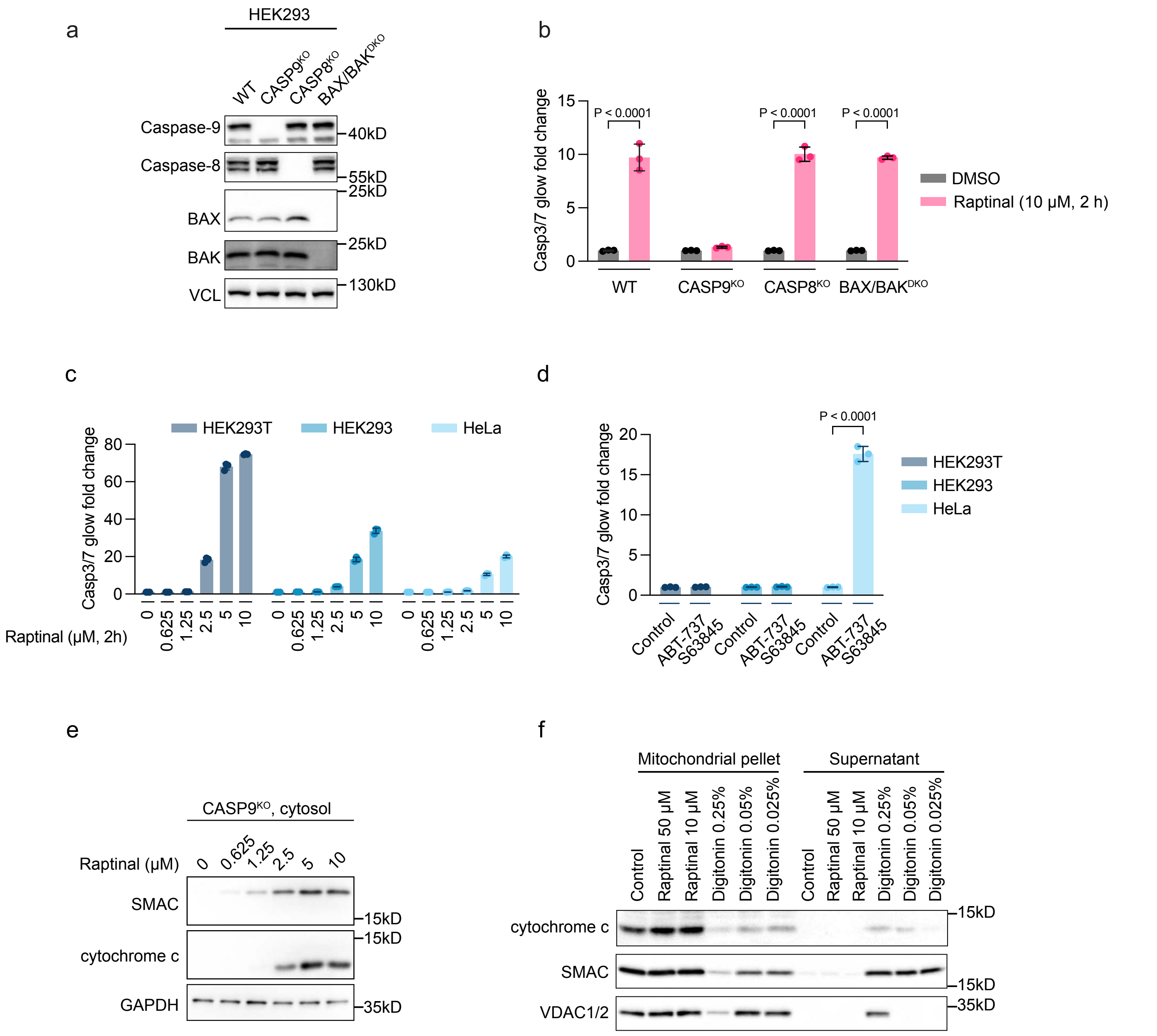


Supplementary Figure 1

**Supplementary Fig. 1 Raptinal induces mitochondrial damage without directly permeabilizing isolated mitochondria**

**(a)** Western blot analysis of caspase-9, caspase-8, BAX, and BAK expression in wild-type, CASP9$^{KO}$, CASP8$^{KO}$, and BAX/BAK$^{DKO}$ HEK293 cells. Vinculin (VCL) was used as a loading control. Representative of 2 independent experiments.

**(b)** Wild-type, CASP9$^{KO}$, CASP8$^{KO}$, and BAX/BAK$^{DKO}$ HEK293 cells were treated with DMSO or Raptinal (10 μM) for 2 h. Caspase-3/7 activity was normalized to control condition of each cell line. Data are mean ± SD; n = 3 independently treated wells, representative of 2 independent experiments. p values were calculated for comparison between DMSO and Raptinal treatment for each genotype by two-way ANOVA with Šídák's correction.

**(c)** HEK293T, HEK293, and HeLa cells were treated with the indicated concentrations of Raptinal for 2 h. Caspase-3/7 activity was measured using Caspase-Glo 3/7. Normalized fold changes in caspase-3/7 activity are shown as mean ± SD. n = 3 independently treated wells, representative of 2 independent experiments.

**(d)** HEK293T, HEK293, and HeLa cells were treated with DMSO or BH3 mimetics, ABT-737 (800 nM) and S63845 (400 nM), for 2 h. Caspase-3/7 activity was normalized to control condition of each cell line. Data are mean ± SD; n = 3 independently treated wells, representative of 2 independent experiments. p values were calculated for comparison between control and BH3 mimetic treatment for each cell line by two-way ANOVA with Šídák's correction.

**(e)** CASP9$^{KO}$ HEK293 cells were treated with the indicated concentrations of Raptinal for 2 h. Cytosolic fractions were extracted by mild digitonin permeabilization and analyzed by western blotting for SMAC and cytochrome c. GAPDH was used as a cytosolic loading control. Representative of 2 independent experiments.

**(f)** Isolated mitochondria were treated with Raptinal at the indicated concentrations or digitonin at the indicated percentages at 37 °C for 30 min. Mitochondrial pellets and supernatants were analyzed by western blotting for cytochrome c, SMAC, and VDAC1/2. Representative of 3 independent experiments.

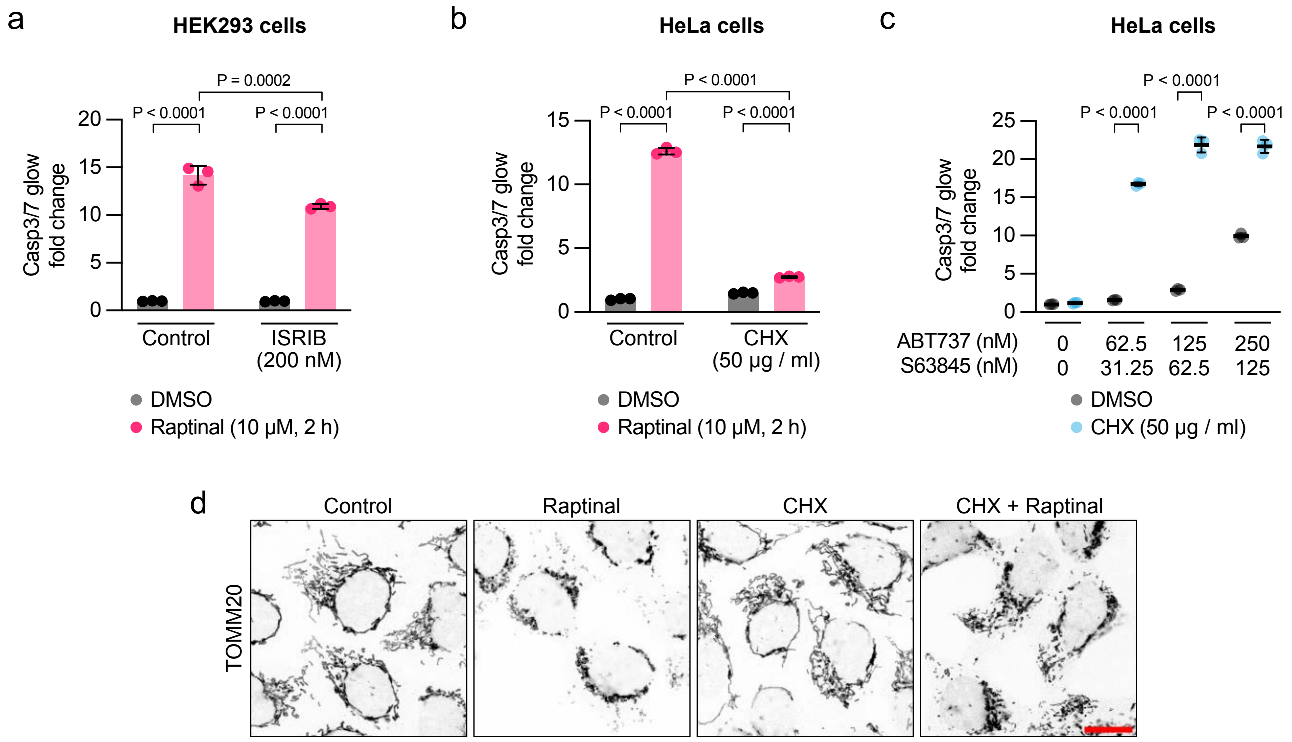


Supplementary Figure 2

**Supplementary Fig. 2 Translation inhibition selectively suppresses Raptinal-induced apoptosis**

**(a)** Wild-type HEK293 cells were pretreated with vehicle or ISRIB (200 nM) for 1 h and then treated with DMSO or Raptinal (10 µM) for 2 h. Caspase-3/7 activity was normalized to DMSO-treated control. Data are mean ± SD; n = 3 independently treated wells, representative of 2 independent experiments; two-way ANOVA with Tukey's correction.

**(b)** HeLa cells were pretreated with vehicle or CHX (50 µg/mL) for 1 h and then treated with DMSO or Raptinal (10 µM) for 2 h. Caspase-3/7 activity was normalized to DMSO-treated control. Data are mean ± SD; n = 3 independently treated wells, representative of 3 independent experiments; two-way ANOVA with Tukey's correction.

**(c)** HeLa cells were pretreated with DMSO or CHX (50 µg/mL) for 1 h and then treated with the indicated concentrations of the BH3 mimetics ABT-737 and S63845 for 2 h. Normalized fold changes in caspase-3/7 activity are shown as mean ± SD. n = 3 independently treated wells, representative of 3 independent experiments. Statistical significance between DMSO and CHX conditions at each BH3 mimetic dose was calculated by two-way ANOVA with Šídák's correction.

**(d)** $CASP9^{KO}$ HEK293 cells were pretreated with vehicle or CHX (50 µg/mL) for 1 h and then treated with DMSO or Raptinal (10 µM) for 2 h. Cells were fixed and immunostained for TOMM20. Representative confocal images are shown. Scale bar, 20 µm.

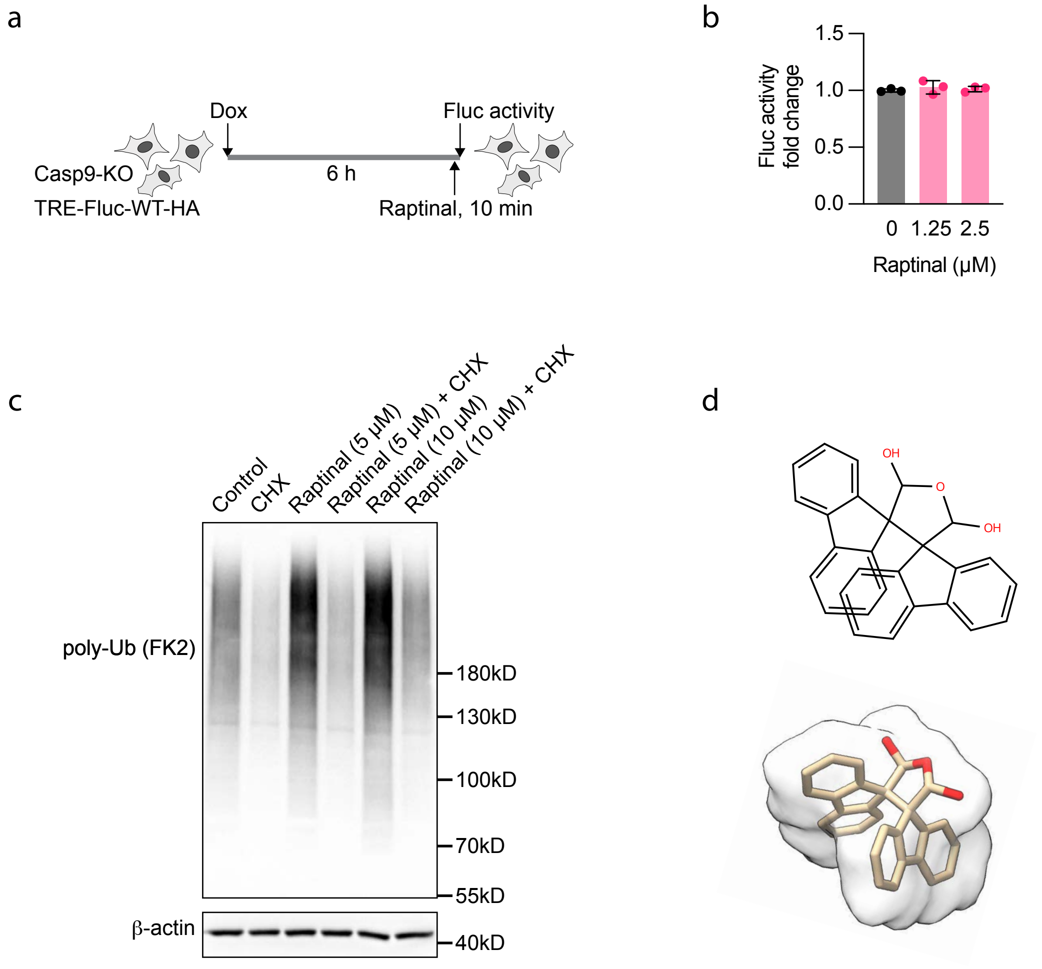


Supplementary Figure 3

**Supplementary Fig. 3 Acute Raptinal exposure does not directly inhibit mature firefly luciferase activity in cells**

**(a)** Experimental design for **(b)**. CASP9$^{KO}$ HEK293 cells inducibly expressing HA-tagged wild-type firefly luciferase (TRE-Fluc-WT-HA) were treated with doxycycline (Dox; 200 ng/mL) for 6 h to induce reporter expression, followed by Raptinal treatment at the indicated concentrations for 10 min before measuring luciferase activity.

**(b)** Firefly luciferase activity from cells treated as in **(a)**. Data are normalized to the control cells without Raptinal treatment and shown as mean ± SD. n = 3 biological replicates.

**(c)** CASP9$^{KO}$ HEK293 cells were pretreated with vehicle or CHX (50 μg/mL) for 1 h and then treated with 5 μM or 10 μM Raptinal for 6 h. Poly-ubiquitinated proteins were probed with anti-ubiquitin antibody (clone FK2). β-actin was used as loading control. Representative of 3 independent experiments.

**(d)** Chemical structure and three-dimensional representation of Raptinal in aqueous solution, showing two fluorenyl groups linked by a cyclic bis-hemiacetal scaffold.

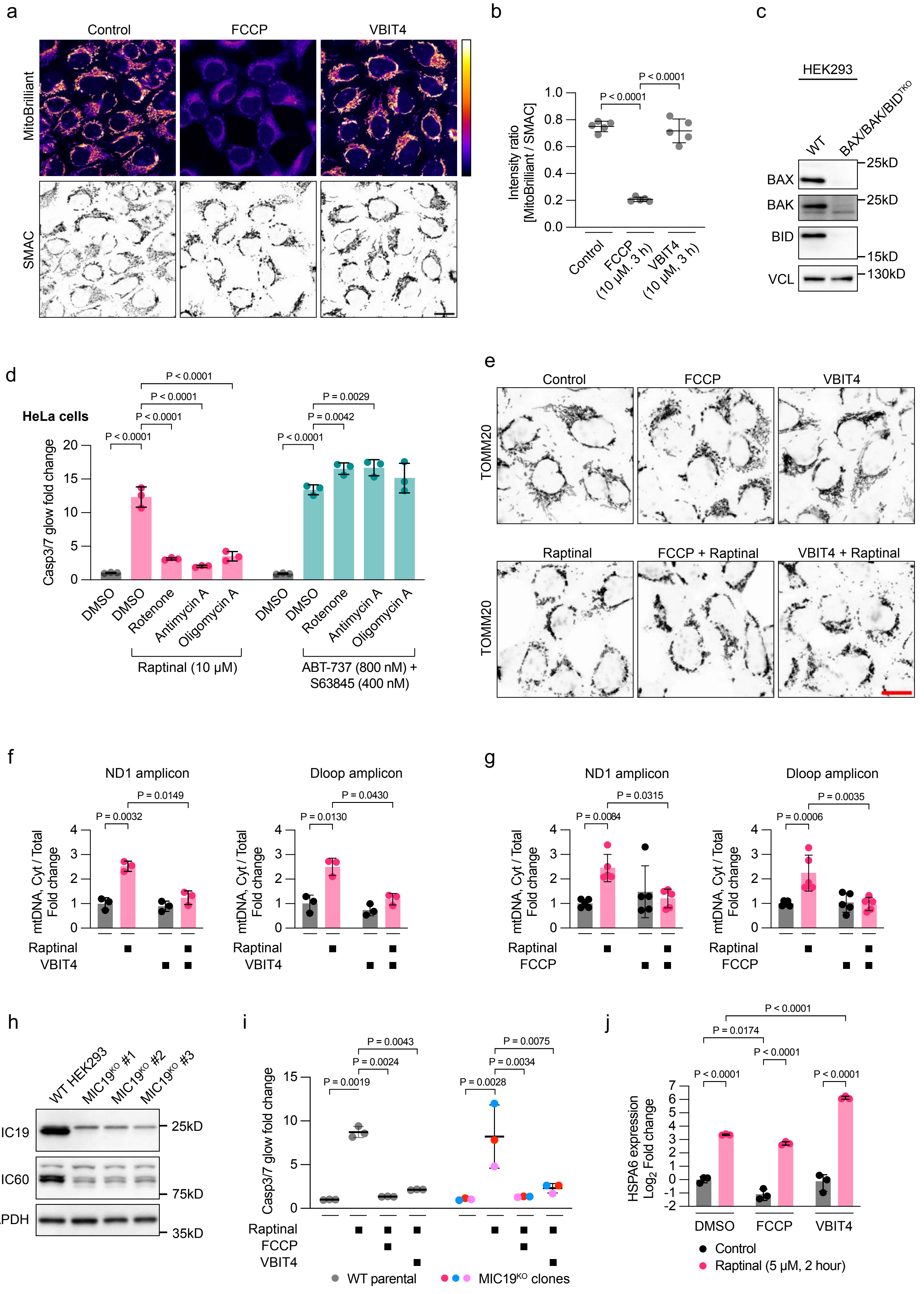


Supplementary Figure 4

**Supplementary Fig. 4 Mitochondrial energetic disruption and VBIT4 uncouple mitochondrial leakage from Raptinal-induced proteotoxic stress**

(**a**) Representative confocal images of wild-type HEK293 cells treated with vehicle, FCCP (10 μM), or VBIT4 (10 μM) for 3 h. Mitochondrial membrane potential was assessed by MitoBrilliant staining, and mitochondria were visualized by SMAC immunostaining. MitoBrilliant signal is shown in Fire LUT. Scale bar, 20 μm.

**(b)** Quantification of MitoBrilliant intensity normalized to SMAC intensity within SMAC-masked mitochondrial area from cells treated as in **(a)**. Each point represents one field; bars indicate mean ± SD; n = 5 fields from 5 independently treated chamber wells; one-way ANOVA with Tukey's correction.

**(c)** Western blot analysis of BAX, BAK, and BID expression in wild-type and BAX/BAK/BID$^{TKO}$ HEK293 cells. VCL, loading control. Representative of 2 independent experiments.

**(d)** HeLa cells were pretreated with vehicle, rotenone (1 μM), antimycin A (10 μM), or oligomycin A (2 μM) for 1 h and then treated with Raptinal (10 μM) or BH3 mimetics, ABT-737 (800 nM) and S63845 (400 nM), for 2 h. Fold changes in caspase-3/7 activity are shown as mean ± SD. n = 3 independently treated wells, representative of 3 independent experiments. Statistical significance relative to cells treated with Raptinal alone or BH3 mimetics alone was calculated by two-way ANOVA with Dunnett's correction.

**(e)** CASP9$^{KO}$ HEK293 cells were pretreated with vehicle, FCCP (10 μM), or VBIT4 (10 μM) for 1 h and then treated with DMSO or Raptinal (10 μM) for 2 h. Cells were fixed and immunostained for TOMM20. Representative confocal images are shown. Scale bar, 20 μm.

**(f)** and **(g)** CASP9$^{KO}$ HEK293 cells were pretreated with vehicle, VBIT4 (2.5 μM) in **(f)**, or FCCP (10 μM) in **(g)** for 1 h and then treated with DMSO or Raptinal (1.25 μM) for 2 h. Cytosolic fractions were extracted by mild digitonin permeabilization, and cytosolic mtDNA was quantified by qPCR using ND1 and Dloop amplicons and normalized to total mtDNA in respective conditions. Data are mean ± SD; n = 3 biological replicates for **(f)** and n = 5 biological replicates for **(g)**. p values were calculated on log-transformed data by two-way ANOVA with Tukey's correction.

**(h)** Western blot analysis of MIC19 and MIC60 expression in wild-type HEK293 cells and three independent MIC19$^{KO}$ clonal cell lines. GAPDH was used as a loading control.

**(i)** Wild-type parental HEK293 cells and MIC19$^{KO}$ clonal cell lines were pretreated with vehicle, FCCP (10 μM), or VBIT4 (10 μM) for 1 h and then treated with DMSO or Raptinal (10 μM) for 2 h. Fold changes in caspase-3/7 activity relative to control condition for each genotype are shown as mean ± SD. n = 3 for wild-type control and 3 MIC19$^{KO}$ clonal cell lines were used as independent biological replicates each individually color-coded. Statistical significance within each genotype was calculated by two-way ANOVA with Tukey's correction.

**(j)** CASP9$^{KO}$ HEK293 cells were pretreated with vehicle, FCCP (10 μM), or VBIT4 (10 μM) for 1 h and then treated with DMSO or Raptinal (5 μM) for 6 h. HSPA6 expression was analyzed by RT-qPCR and is shown as fold change after normalization to GAPDH. Data are mean ± SD; n = 3 biological replicates. p values were calculated on log-transformed data by two-way ANOVA with Tukey's correction.

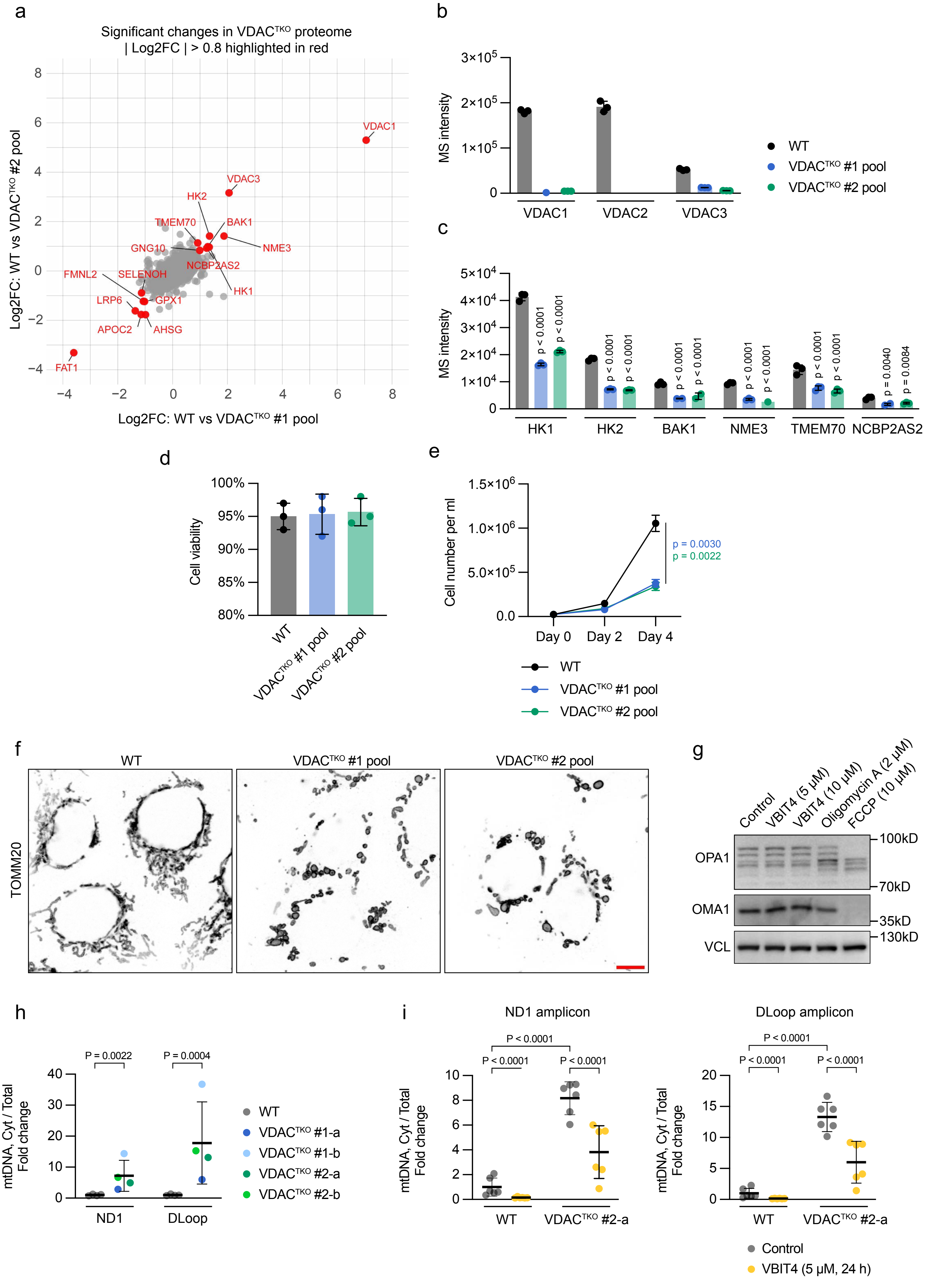


Supplementary Figure 5

**Supplementary Fig. 5 Characterization of VDAC-deficient cells and mitochondrial DNA leakage**

**(a)** Proteomic comparison between wild-type HEK293 cells and two pooled VDAC1/2/3$^{TKO}$ cell populations. Scatter plot shows $log_2$ fold change for proteins in wild-type cells relative to VDAC1/2/3$^{TKO}$ pool #1 and pool #2. Proteins meeting nominal $p < 0.05$ and $|log_2$ fold change$| > 0.8$ are highlighted in red. n = 3 biological replicates per condition.

**(b)** Raw MS intensities of VDAC1, VDAC2, and VDAC3 in wild-type HEK293 cells and two pooled VDAC1/2/3$^{TKO}$ cell populations. Missing values are not imputed. Data are mean ± SD; n = 3 biological replicates per condition.

**(c)** Raw MS intensities of selected mitochondrial proteins reduced in pooled VDAC1/2/3$^{TKO}$ cells. Missing values are not imputed. Data are mean ± SD; n = 3 biological replicates per condition. Statistical significance relative to wild-type cells for each protein was calculated by two-way ANOVA with Dunnett's correction.

**(d)** Cell viability of wild-type HEK293 cells and two pooled VDAC1/2/3$^{TKO}$ cell populations under basal culture conditions was assessed by trypan blue staining. Data are mean ± SD. n = 3 biological replicates.

**(e)** Proliferation of wild-type HEK293 cells and two pooled VDAC1/2/3$^{TKO}$ cell populations over 4 days. Cell numbers are shown as mean ± SD. n = 3 biological replicates. p values comparing each VDAC1/2/3$^{TKO}$ pool with wild-type cells at day 4 were calculated by two-way ANOVA with Dunnett's correction.

**(f)** Representative confocal images of wild-type HEK293 cells and two pooled VDAC1/2/3$^{TKO}$ cell populations immunostained for TOMM20. Scale bar, 10 μm.

**(g)** CASP9$^{KO}$ cells were treated with VBIT4, oligomycin A and FCCP at indicated concentrations for 8 h. OPA1 processing and OMA1 expression were analyzed by western blotting. VCL, loading control. Representative of 3 independent experiments.

**(h)** Cytosolic mtDNA in wild-type HEK293 cells and 4 independent VDAC1/2/3$^{TKO}$ clonal cell lines. Cytosolic fractions were extracted by mild digitonin permeabilization, and cytosolic mtDNA was quantified by qPCR using ND1 and Dloop amplicons and normalized to total mtDNA in respective conditions. Ratios between cytosolic mtDNA and total mtDNA are further normalized to that of wild-type cells under control condition and shown as mean ± SD. n = 4 biological replicates for wild-type cells and each VDAC1/2/3$^{TKO}$ clonal cell line was used as an independent biological replicate. Statistical significance relative to wild-type cells for each amplicon was calculated on log-transformed data by unpaired Student's t-test.

**(i)** Wild-type HEK293 cells and VDAC1/2/3$^{TKO}$ clone #2-a were treated with vehicle or VBIT4 (5 μM) for 24 h. Cytosolic mtDNA was quantified as in **(h)**. Data are mean ± SD; n = 6 biological replicates. p values were calculated on log-transformed data by two-way ANOVA with Tukey's correction.

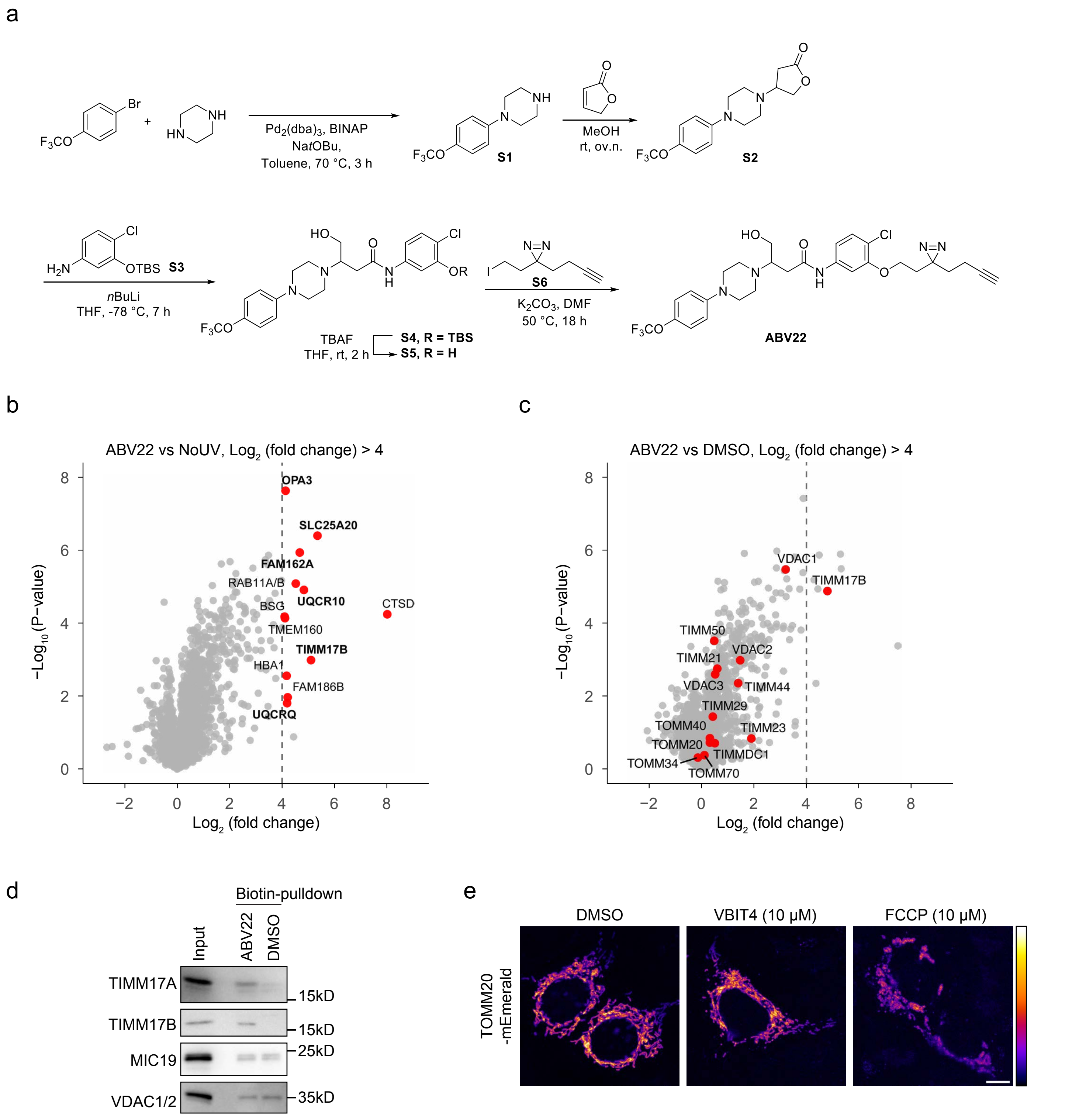


Supplementary Figure 6

**Supplementary Fig. 6 Synthesis and proteomic profiling of the photoreactive VBIT4 analog ABV22**

**(a)** Synthetic route for the photoreactive clickable VBIT4 analog ABV22.

**(b)** Volcano plot showing proteins enriched in ABV22-crosslinked samples compared with samples without UV irradiation. Selected proteins with $\log_2$ fold change > 4 are highlighted in red and mitochondrial proteins are labeled in bold. n = 4 biological replicates per condition. p values were calculated from $\log_2$-transformed LFQ intensities using Welch's t test.

**(c)** Volcano plot showing proteins enriched in ABV22-crosslinked samples compared with DMSO-treated controls. Mitochondrial import machineries and VDAC channels are annotated. n = 4 biological replicates per condition. p values were calculated from $\log_2$-transformed LFQ intensities using Welch's t test.

**(d)** Selective labeling of TIMM17A/B by ABV22. HEK293 cells were incubated with or without ABV22 (1 μM) for 1 h, followed by UV photocrosslinking, click chemistry with biotin-azide, streptavidin pulldown, and immunoblotting for TIMM17A, TIMM17B, VDAC1/2, and MIC19. Input represents total lysate before pulldown. Representative of 2 independent experiments.

**(e)** Representative confocal images of HEK293 cells transiently expressing TOMM20-mEmerald and treated with DMSO, VBIT4 (10 μM), or FCCP (10 μM) for 6 h. TOMM20-mEmerald signal is shown in Fire LUT. Scale bar, 10 μm.

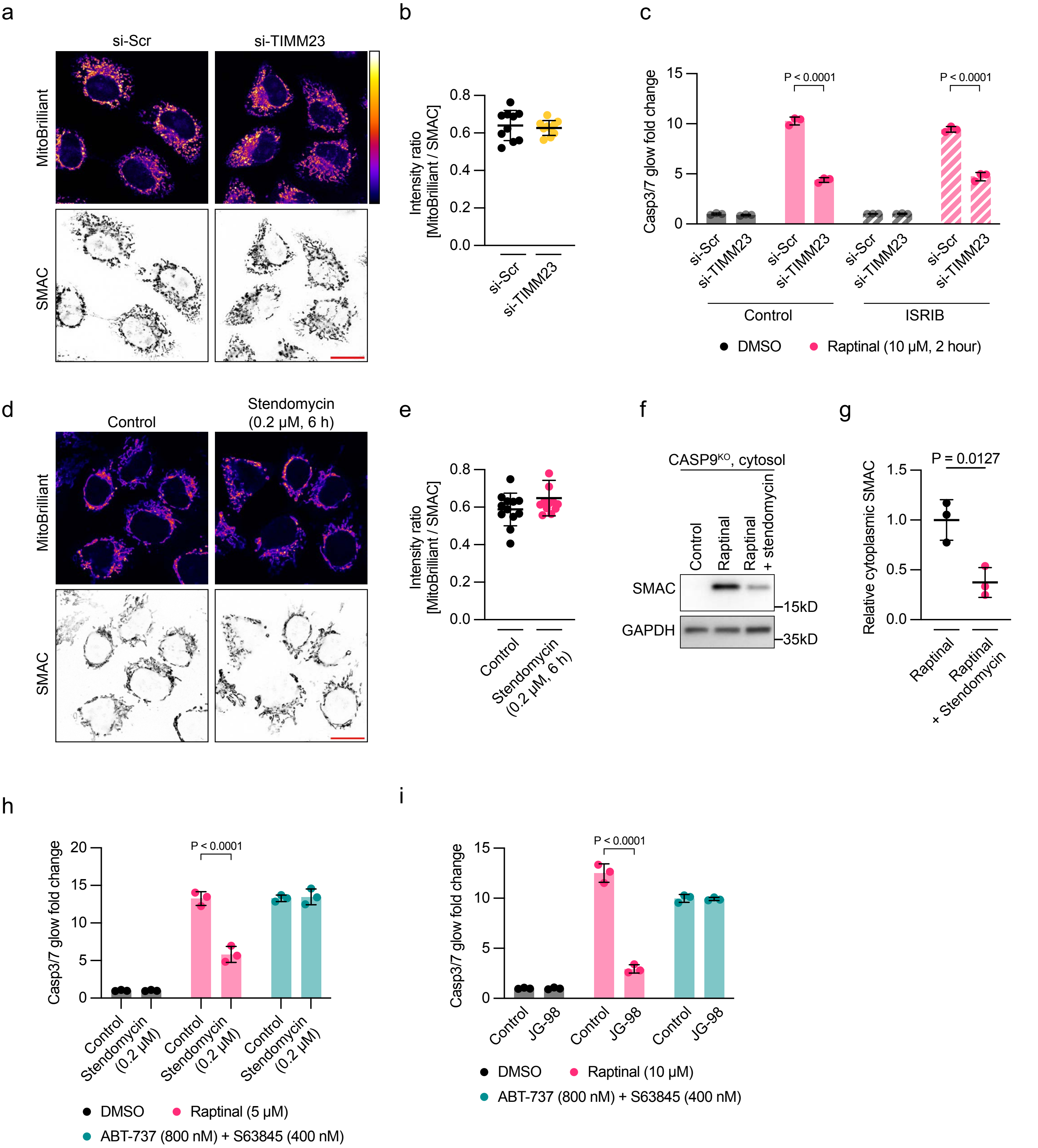


Supplementary Figure 7

**Supplementary Fig. 7 Restriction of the TIM23–PAM axis protects against Raptinal-induced mitochondrial damage**

**(a)** HeLa cells were transfected with non-targeting control siRNA (si-Scr) or TIMM23-targeting siRNA (si-TIMM23) for 48 h. Mitochondrial membrane potential was assessed by MitoBrilliant staining, and mitochondria were visualized by SMAC immunostaining. Representative confocal images are shown. Scale bar, 20 μm.

**(b)** Quantification of MitoBrilliant intensity normalized to SMAC intensity within SMAC-masked mitochondrial areas from cells treated as in **(a)**. Each point represents one field; n = 10 fields per condition pooled from 3 independent transfections; bars indicate mean ± SD.

**(c)** HEK293 cells were transfected with si-Scr or si-TIMM23 in the presence or absence of ISRIB (200 nM) for 48 h and then treated with DMSO or Raptinal (10 μM) for 2 h. Fold changes in caspase-3/7 activity relative to DMSO-treated control condition are shown as mean ± SD. n = 3 independently treated wells, representative of 2 independent experiments. Statistical significance between si-Scr- and si-TIMM23-transfected cells under each condition was calculated by two-way ANOVA with Šídák's correction.

**(d)** HEK293 cells were pretreated with vehicle or stendomycin (200 nM) for 5 h. Mitochondrial membrane potential was assessed by MitoBrilliant staining, and mitochondria were visualized by SMAC immunostaining. Representative confocal images are shown. Scale bar, 20 μm.

**(e)** Quantification of MitoBrilliant intensity normalized to SMAC intensity within SMAC-masked mitochondrial areas from cells treated as in **(d)**. Each point represents one field; n = 12 fields per condition pooled from 3 independently treated chamber wells; bars indicate mean ± SD.

**(f)** $CASP9^{KO}$ HEK293 cells were pretreated with vehicle or stendomycin (200 nM) for 5 h and then treated with DMSO or Raptinal (5 μM) for 2 h. Cytosolic fractions were extracted by mild digitonin permeabilization and analyzed by western blotting for SMAC. GAPDH was used as a cytosolic loading control. Representative of 3 independent experiments.

**(g)** Quantification of cytosolic SMAC from cells treated as in **(f)**, expressed as relative change over Raptinal-treated samples. Data are mean ± SD; n = 3 independent experiments. Statistical significance was calculated using unpaired Student's t-test.

**(h)** HeLa cells were pretreated with vehicle or stendomycin (200 nM) for 24 h and then treated with DMSO, Raptinal (10 μM), or BH3 mimetics, ABT-737 (800 nM) and S63845 (400 nM), for 2 h. Caspase-3/7 activity was normalized to DMSO-treated control. Data are mean ± SD; n = 3 independently treated wells, representative of 2 independent experiments. Statistical significance between vehicle- and stendomycin-pretreated cells within each treatment condition was calculated by two-way ANOVA with Šídák's correction.

**(i)** HeLa cells were pretreated with vehicle or JG-98 (2 μM) for 1 h and then treated with DMSO, Raptinal (10 μM), or BH3 mimetics, ABT-737 (800 nM) and S63845 (400 nM), for 2 h. Fold changes in caspase-3/7 activity relative to DMSO-treated control condition are shown as mean ± SD. n = 3 independently treated wells, representative of 2 independent experiments. Statistical significance between vehicle- and JG-98-pretreated cells within each treatment condition was calculated by two-way ANOVA with Šídák's correction.

### Supplementary Tables

**Supplementary Table 1. DESeq2 analysis of Raptinal-treated cells, related to Figure 1**

RNA-seq differential expression analysis of DMSO- and Raptinal-treated CASP9$^{KO}$ HEK293 cells, including normalized counts, $log_2$ fold changes, Wald statistics, p values, adjusted p values, gene identifiers, symbols, descriptions, and biotypes.

**Supplementary Table 2. Whole-cell proteomic profiling of VDAC$^{TKO}$ cells, related to Figure 4 and Supplementary Fig. 5**

Whole-cell proteomic profiling of wild-type HEK293 cells and two pooled VDAC$^{TKO}$ cell populations. Raw MS intensities are shown for three biological replicates per condition. Missing values were not imputed.

**Supplementary Table 3. ABV22 chemical proteomics, related to Figure 5 and Supplementary Fig. 6**

LC-MS/MS analysis of ABV22-enriched proteins from HEK293 cells treated with DMSO, ABV22 plus UV photocrosslinking, or ABV22 without UV irradiation. The table includes LFQ intensities from four biological replicates per condition, MitoCarta annotation, $log_2$ fold changes, and nominal p values calculated from $log_2$-transformed LFQ intensities for ABV22-crosslinked samples relative to both controls.

**Supplementary Table 4. High-enrichment ABV22-associated proteins, related to Figures 5 and Supplementary Fig. 6**

Subset of ABV22-associated proteins with $log_2$ fold change > 3 and nominal p value < 0.05 relative to both DMSO and ABV22 without UV controls. The table includes protein identifiers, enrichment statistics, MitoCarta annotation, transmembrane association, and subcellular localization.

**Supplementary Table 5. Antibodies, chemical tools and oligonucleotides used in this study**
